# OpenTree: A Python package for Accessing and Analyzing data from the Open Tree of Life

**DOI:** 10.1101/2020.12.14.422759

**Authors:** Emily Jane McTavish, Luna Luisa Sanchez Reyes, Mark T. Holder

## Abstract

The Open Tree of Life project constructs a comprehensive, dynamic and digitally-available tree of life by synthesizing published phylogenetic trees along with taxonomic data. Open Tree of Life provides web-service application programming interfaces (APIs) to make the tree estimate, unified taxonomy, and input phylogenetic data available to anyone. Here, we describe the python package `opentree`, which provides a user friendly python wrapper for these APIs and a set of scripts and tutorials for straightforward downstream data analyses. We demonstrate the utility of these tools by generating an estimate of the phylogenetic relationships of all bird families, and by capturing a phylogenetic estimate for all taxa observed at the University of California Merced Vernal Pools and Grassland Reserve.

---

Evolutionary history provides a framework to link all life on earth. However, it is not easy to access accurate, up-to-date phylogenetic relationships for arbitrary sets of taxa of interest, even if phylogenetic estimates for those taxa have been made and published [Drew et al., 2013, McTavish et al., 2017]. Individual phylogenetic estimates are not comprehensive, and therefore seldom contain all taxa of interest. Taxonomic relationships, while comprehensive, provide a coarse, and often outdated, picture of shared ancestry. The Open Tree of Life project (`opentree`) provides a reproducible framework for accessing up-to-date evolutionary relationships for arbitrary sets of taxa across the entire tree of life. All data in `opentree` is freely available via API’s. The package `opentree` provides a user friendly python interface to access these data. In addition `opentree` is packaged with a set of tutorials and scripts to make common downstream analyses straightforward.

## Description

`opentree` is a Python package for accessing and analyzing data from the openTree of Life project. Open Tree of Life stores a wealth of taxonomic and phylogenetic data gathered together in an open-access interoperable framework. The current synthetic tree [openTree Of Life et al., 2019] comprises 2.4 million tips. Most of the tips of the tree represent species, but some are infraspecific taxa. The framework of this tree is provided by a unified taxonomy [OpenTreeofLife et al., 2019, Rees and Cranston, 2017]. This taxonomy links unique identifiers across many online taxonomic resources, including NCBI [Federhen, 2012], GBIF [GBIF, 2019], as well as user contributed taxonomic amendments contained in [https://github.com/OpenTreeOfLife/amendments-1]. These taxonomic relationships are refined by evolutionary estimates from 1,216 published papers including 87,000 tip taxa [OpenTreeOfLife et al., 2019, Redelings and Holder, 2017]. The Open Tree data store, ‘Phylesystem’ [McTavish et al., 2015] contains 4,500 published studies, including those incorporated in the tree. Each tree has mappings between the tips in these published studies and unique taxonomic identifiers.

All of these data are freely accessible via API calls, documented at https://github.com/OpenTreeOfLife/germinator/wiki/Open-Tree-of-Life-Web-APIs. `opentree` provides an user-friendly wrapper for calling these APIs. In addition, it converts these data between commonly used file formats and data types. This package allows allows users to generate to data objects in DendroPy, a phylogenetic computing library [Sukumaran and Holder, 2010].

`opentree` incorporates in python the functionality available in rotl, an R package to interact with the Open Tree of Life data [Michonneau et al., 2016], as well as additional downstream analysis and interoperability tools. rotl has been already been cited 132 times in the 4 years since its publication, demonstrating a demand for accessible user access to these data. By providing a python package to interact with these data, we make it straightforward for python users to access and analyze these data. A python wrapper for Open Tree of Life also makes linking these data with the stable of other Python biodiversity informatics tools straightforward.

In addition, `opentree` expands the toolset available for working with the `opentree` unified taxonomy [Rees and Cranston, 2017].

## Services Provided by Opentree

The openTree APIs are divided into three main categories, synthetic tree, taxonomy and taxonomic name resolution, and study search. Many analyses integrate calls to each of these subcomponents. The `opentree` package links across these services to make common API calls easier. Some example calls are described here, but all methods and scripts are fully documented, including examples and return formats at https://opentree.readthedocs.io.

### Synthetic tree

The openTree synthetic tree contains all 2.4 million taxa in the openTree taxonomy, with relationships for 87,000 taxa informed by 1,216 studies. Each branch in the tree is informed by published phylogenetic relationships, where they are present in the curated data store, or by taxonomic relationships where no phylogenetic data is available. For each node in the synthetic tree, the API returns identifiers for all the trees in the synthesis pipeline which support or conflict with that node. Each node is uniquely labeled. If the descendants of a node align with named taxonomic group, the taxon identifier is applied to the node. If the node does not correspond a taxon named in the openTree taxonomy, the node is labeled using a phyloreference [Parr et al., 2012] describing that node as the most recent common ancestor of two identified taxa. `opentree` users can easily access evolutionary estimates for arbitrary sets of taxa. The web-service response also includes the published phylogenetics estimates which underlie those inferences. The `opentree` wrapper captures and formats these citations to make providing appropriate credit for these synthetic induced subtree estimates straightforward. Users can also access full synthetic subtrees subtending any individual node.

### Taxonomy and Taxonomic Name Resolution

The openTree taxonomy not only provides a scaffold for the synthetic tree, but is also a valuable resource in its own right. Matching names is a key hurdle in bioinformatics. Correct taxon names change through time, and spelling and typographical errors can propagate through bioinformatic resources. Thus, demanding exact matching of names from different sources can be too stringent and fail to match the same taxon. However, different names can be very close in spelling to one another. So, tolerating misspellings makes it easy to accidentally match names that should refer to two distinct taxa. The openTree taxonomy [Rees and Cranston, 2017, OpenTreeofLife et al., 2019] provides a link between the unique identifiers generated by several large scale online taxonomic resources [GBIF, NCBI, Silva, Worms], as well as all known name synonomies provided by those resources. The `opentree` taxonomic name resolution service (TNRS) searches this full taxonomy and returns exact or fuzzy matches from input names string to taxa and their unique identifiers. This TNRS forms a link between human readable name strings, and rigorous unique identifiers.

`openTree` wraps the openTree taxonomy and TNRS APIs for ease of integrating taxonomy and TNRS queries with downstream analyses. In addition, `opentree` provides helpers for quickly searching the text download of the taxonomy, which can be more efficient for bulk queries.

### Study search

The OpenTree datastore contains 4,468 published phylogenetic studies, including 9,395 phylogenetic trees (as of Dec 4, 2020). These studies and trees are indexed on a number of properties including author name, curator name, and publication information. In addition, the tips of these trees are mapped to the unified Open Tree taxonomy making comparisons among estimates of relationships and searching for taxa of interest straightforward. This allows searching of studies based on taxa contained in the study. Importantly, this search does not rely on string-matching of what the taxonomic name was at the time of publication - rather it leverages the full suite of synonomies gathered across the input taxonomies to find equivalent taxa across studies, even if the canonical name has changed between publications. The indexing of these trees is taxonomically explicit. So, for example, a search for ‘canidae’ will find trees with taxa contained in the taxonomic group *canidae*, even if the term ‘canidae’ does not itself appear in the tree or tips. Based on the results of these searches, studies can be viewed in a browser on the OpenTree curator site, or the phylogenies themselves can be downloaded for comparisons or other downstream use.

In addition, as the tips of each study are mapped by curators to identifiers in the OpenTree taxonomy, comparing the relationships represented in input trees to taxonomic relationships and to taxonomy is straightforward. The browser based tree viewer has a graphical visualization of this concordance and conflict. `opentree` provides a wrapper for this conflict functionality, which makes it straightforward to assess what taxon definitions and evolutionary relationships a tree estimate agrees with and conflicts with. This functionality can also be applied to local phylogenies for which the tips have been matched to taxonomy. This allows users to assess concordance and conflict with previous inferences in pre-publication trees, even without sharing them to the publicly available OpenTree data store [Reyes et al., 2020, McTavish et al., 2015].

## Biological Examples

There are a plethora of downstream applications of this linked set of resources. We highlight two examples based on user queries.

### A phylogeny of all bird families

A full Jupyter notebook tutorial demonstrating how to access a tree of all bird families is packaged with the software at https://github.com/OpenTreeOfLife/python-opentree/blob/main/docs/notebooks/TreeOfBirdGenera.ipynb. Capturing evolutionary information at large scales is often simplified by using arbitrary taxonomic cutoffs. While the OpenTree taxonomy is not rank focused, it does track rank information from component taxonomies. By searching the OpenTree taxonomy for families in birds, we find that there are 390 listed bird families, 196 of which are included in the synthetic tree. Groups that are excluded from the synthetic tree for a few potential reasons, the most common of which is that all members of the group are extinct, and we have no curated published studies providing information about the correct evolutionary relationships. Placements of fossil taxa based only on taxonomy tend to be unstable, and the OpenTree synthesis procedure excludes taxa if the taxon is not present in at least one phylogenetic input. These families can be included in later synthetic trees if data is added to the study corpus providing information on their relationships. Other taxa are excluded from synthesis if issues have been raised about their taxonomic validity, such as if the name corresponds to a family that is ‘barren’, i.e. it contains no species in the OpenTree taxonomy, or because the name was judged to be invalid by the OpenTree taxonomy merging software[Rees and Cranston, 2017].

If we request an induced subtree from the synthesis for these 164 taxa, we get back a tree that has 150 tips. The return value also includes a list of non-monophyletic taxa. Some of the non-monophyletic taxa map to internal nodes on our output tree. In those cases, input phylogenies are telling us that these ‘families’ are paraphyletic with respect to other families. Which studies contest the monophyly of a taxonomic clade can be easily accessed through the browser (e.g. https://tree.opentreeoflife.org/opentree/argus/ottol@603925) or via the `opentree` wrapper using queries to `opentree.synth subtree`. Figure 1 shows the shows the topology of the 130 monophyletic bird families plus MRCA’s of 20 additional non-monophyletic families as tips. The other 14 taxa are non-monophyletic families for which the MRCA is an internal node on the output tree.

**Fig. 1.**
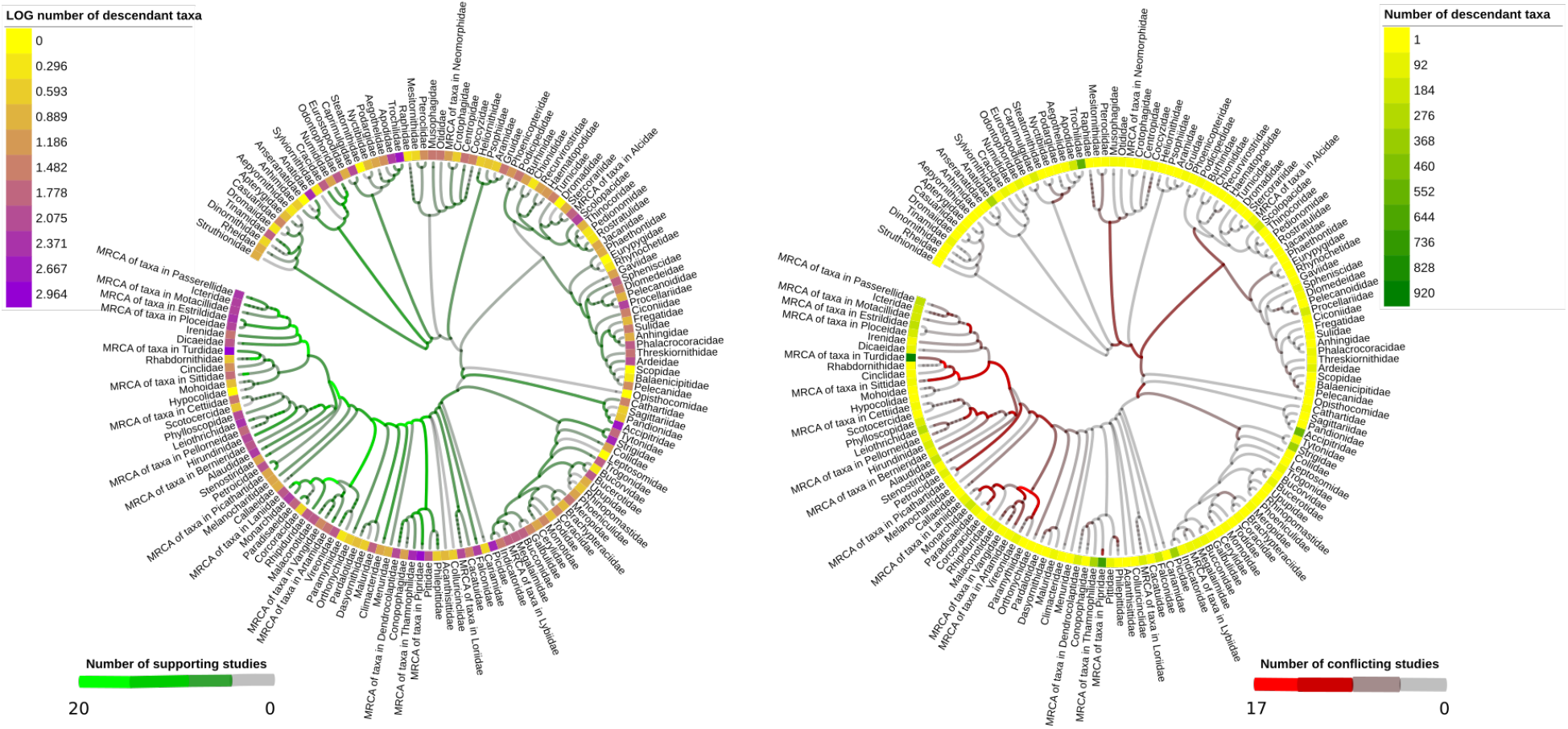
Relationships of 150 bird families, showing the number of published phylogenetic studies which support and and conflict with each branch in the tree. For families which are not monophyletic according to published phylogenies, tips for those families are labelled with ‘MRCA of taxa in Family name’. Heat maps show the number of tip taxa descendants from each tip, in log scale on left, and in exact numbers on left. Branch colors show the number of input studies which support (left, green) or conflict with (right, red) each inferred branch in the super tree. Branch lengths are arbitrary. A total of 64 published phylogenies underlie the relationships in this tree (Citations in supplemental information). Figure created using [Letunic and Bork, 2019]

These families contain from 1 to 920 total descendant tip taxa (species and subspecies). Across all 150 families, 10,357 descendent tip taxa are captured by the relationships shown in this tree. Figure 1 displays the number of descendant taxa in each family as a heat map, with log of the number of descendants displayed on the left, and the actual number of descendents on the right. This display demonstrates that ‘families’ is not a very even way to break up biodiversity across birds.

Based on the phylogenetic synthesis algorithm, each branch in the tree is supported by either taxonomy alone, where there are no input studies that traverse that branch, or by one or more input phylogenetic studies. The support for each node in the tree can be interrogated using an `opentree.synth node` info call. While each branch must be supported by taxonomy or at least one input study from phylesystem, where multiple inputs traverse a branch, there can be conflict among studies. The synthesis algorithm is greedy, and the output tree will display the branch supported by the highest ranked study included in synthesis. The node info call will return not only which studies support that branch, but also which studies have relationships which conflict with that branch. In figure 1 support or conflict for each branch is displayed by the intensity of green and red coloration, respectively. Some branches in this tree are supported by 20 studies, and a few show conflict with up to 17 other studies. Of the 443 branches in this tree, 422 are supported by at least one input phylogenetic study, and the other 21 are based on taxonomic relationships.

This tree shows only topology. When combining taxonomy, and phylogenetic branches from across studies with vastly different data types, merging branch lengths is not meaningful. However, estimates of node ages can be gathered using downstream tools such as DateLife [Sánchez-Reyes and O’Meara, 2019] which gathers and synthesizes node date information from studies in the phylesystem corpus.

### Linking data from the Global Biodiversity Information Facility (GBIF) with phylogenetic information from Open Tree of Life

The University of California, Merced has a natural reserve directly adjacent to campus, which contains several vernal pools. These vernal pools create a unique habitat which allows native species to thrive, and the proximity to campus allows undergraduate classes to experience this ecosystem on field trips which can be accomplished during class time. A species list for the reserve and adjacent campus areas is available through Global Biodiversity Information Facility (GBIF) website [GBIF, 2019]. GBIF provides a repository for species occurrence data tracked in a variety of data stores, including bird observations from eBird [Sullivan et al., 2009], community science observations from iNaturalist (www.inaturalist.org), and several other resources. A full tutorial demonstrating how to access a tree for a GBIF data download is packaged with the software at https://github.com/OpenTreeOfLife/python-opentree/blob/main/docs/notebooks/gbif/GBIF_to_OpenTree.ipynb.

We downloaded the full list of animal observations from the UC Merced Vernal pools reserve from GBIF [GBIF, 2019]. This data download comprised 6,709 records from 223 species. Using the GBIF unique taxon identifiers, 201 of these species could be directly matched to taxa in the OpenTree taxonomy using opentree.taxon info(source id = gbif unique identifier). This direct matching captures exact one to one relationships between these taxonomies, and avoids slow and potentially error prone string matching. Nineteen taxa had updated identifiers in GBIF since the most recent reconciliation between the GBIF taxonomy and the OpenTree taxonomy, and were assigned OpenTree taxon identifiers based on exact string matches. There were two taxa “*P. abortivum* St.” and “*Ichneumon cupitus* Cresson 1877”, which were not found in the OpenTree taxonomy, and were dropped from the analysis.

Using this set of 223 OpenTree unique identifiers, an induced synthetic tree for these taxa can be downloaded (Figure 2). This synthetic tree is supported by 160 individual published trees (Citations in supplemental information)).

**Fig. 2.**
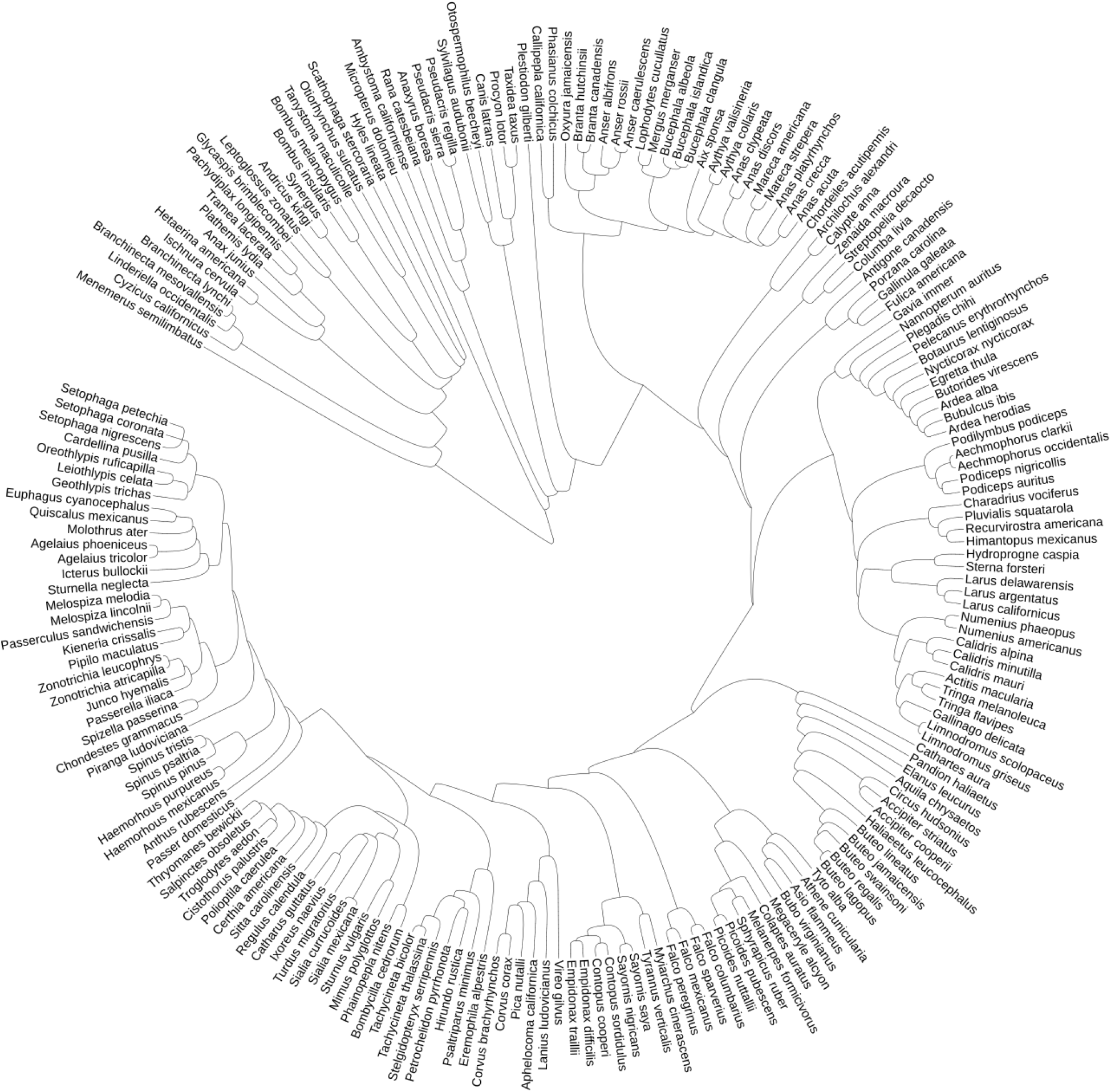
Evolutionary relationships between all animal taxon records in the UC Merced Vernal Pools and Grassland Reserve. Branch lengths are arbitrary. A total of 160 published phylogenies underlie the relationships in this tree (Citations in supplemental information). Figure created using [Letunic and Bork, 2019]

For researchers, working in the vernal pools reserve, this tree also provides the necessary information for community phylogenetic analyses. Li et al. [2019] demonstrated that synthetic phylogenies from the OpenTree project perform well in community phylogenetic studies. By providing ready access to these estimates, based on 160 previously published trees, opentree makes basing ecological analyses in an accurate evolutionary framework straightforward.

The ability to build a phylogeny of local taxa is also a valuable pedagogical tool. One of us (EJM) used this phylogeny to discuss the diversity of life of animal life as part of a class exercise on vernal pools ecology and evolution, in an undergraduate evolution class. Students visited the UC Merced Vernal Pools and Grassland Reserve, and then explored the evolutionary tree of all the animal species recorded as observed in the reserve. There are several threatened and endangered species on the vernal pools reserve, including two species of fairy shrimp, *Branchinecta lynchii* (threatened) and *Branchinecta mesovallensis* (endangered). By building a phylogenetic tree of taxa found on and around campus, tree thinking examples in class can have a direct connection for students. For example, this tree (Figure 2) shows that the genus *Anas* does not form a monophyletic group. Walking the tree of life has been demonstrated to be an effective way to get students to understand the connections among different lineages of life on earth [Ballen and Greene, 2017]. Walking through this tree, and labelling major animal groups allows students to connect to the diversity of animal life based on the actual species surrounding them, rather than arbitrary textbook examples.

## Supporting information

Supplemental Citations, Vernal Pools tree

Supplemental Citations, Bird Families tree

## Availability

OpenTree is fully open source with a CC0 license. It is available on GitHub https://github.com/OpenTreeOfLife/python-opentree. It can be installed from PyPi using pip install opentree. The code is packaged with an automated test suite which is maintained to cover at minimum 75% of the code. Testing reports are generated on travis.io and posted to codecov.io and reflected on the GitHub readme with each commit. Documentation and tutorials are available with the code, and is posted to https://opentree.readthedocs.io.

## Acknowledgements

This package relies on the OpenTree API’s and the team that continues to develop and improve them, currently Jim Allman, Karen Cranston, Ben Redelings, and the authors of this package. Funding for this project was provided by NSF ABI 1759846 and NSF ABI 1759838. We acknowledge logistical support provided by the UC Merced Vernal Pools and Grassland Reserve. Thank you to Emily Sessa and the organizers of the Society of Systematic Biologists SSB 2020 meeting for supporting the workshop where we tested and demonstrated this package.

