## Supplemental Citations, Vernal Pools tree for "OpenTree: A Python package for Accessing and Analyzing data from the Open Tree of Life"

Citations for the phylogenies underlying the Merced Vernal Pools Animal tree

https://tree.opentreeoflife.org/curator/study/view/ot_816?tab=trees&tree=tree1
Gibson, Rosemary, Allan Baker. 2012. Multiple gene sequences resolve phylogenetic relationships in the shorebird suborder Scolopaci (Aves: Charadriiformes). Molecular Phylogenetics and Evolution 64 (1): 66-72
http://dx.doi.org/10.1016/j.ympev.2012.03.008

https://tree.opentreeoflife.org/curator/study/view/ot_1268?tab=trees&tree=tree5
Martín J. Ramírez, 2014, 'The Morphology And Phylogeny Of Dionychan Spiders (Araneae: Araneomorphae)', Bulletin of the American Museum of Natural History, vol. 390, pp. 1-374
http://dx.doi.org/10.1206/821.1

https://tree.opentreeoflife.org/curator/study/view/pg_1776?tab=trees&tree=tree3581
Cho, S., Zwick A., Regier J., Mitter C., Cummings M.P., Yao J., Du Z., Zhao H., Kawahara A.Y., Weller S.J., Davis D.R., Baixeras J., Brown J.W., & Parr C. 2011. Can deliberately incomplete gene sample augmentation improve a phylogeny estimate for the advanced moths and butterflies (Hexapoda: Lepidoptera)?. Systematic Biology 60 (6): 782-796.
http://dx.doi.org/10.1093/sysbio/syr079

https://tree.opentreeoflife.org/curator/study/view/pg_2664?tab=trees&tree=tree6201
Regier, JC, Mitter C, Zwick A, Bazinet AL, Cummings MP, et al. (2013) A large-scale, higher-level, molecular phylogenetic study of the insect order Lepidoptera (moths and butterflies). PLoS ONE 8 (3): e58568
http://dx.doi.org/10.1371/journal.pone.0058568

https://tree.opentreeoflife.org/curator/study/view/ot_425?tab=trees&tree=tree1
Stein, R. Will, Joseph W. Brown, Arne Ø. Mooers. 2015. A molecular genetic time scale demonstrates Cretaceous origins and multiple diversification rate shifts within the order Galliformes (Aves). Molecular Phylogenetics and Evolution 92: 155-164
http://dx.doi.org/10.1016/j.ympev.2015.06.005

https://tree.opentreeoflife.org/curator/study/view/ot_764?tab=trees&tree=tree1
EMMANUEL F. A. TOUSSAINT, MATTHIAS SEIDEL, EMMANUEL ARRIAGA-VARELA, JIŘÍ HÁJEK, DAVID KRÁL, LUKAŠ SEKERKA, ANDREW E. Z. SHORT, MARTIN FIKÁČEK, 2016, 'The peril of dating beetles', Systematic Entomology
http://dx.doi.org/10.1111/syen.12198

https://tree.opentreeoflife.org/curator/study/view/ot_809?tab=trees&tree=tree2
Jetz, W., G. H. Thomas, J. B. Joy, K. Hartmann, A. O. Mooers. 2012. The global diversity of birds in space and time. Nature 491 (7424): 444-448
http://dx.doi.org/10.1038/nature11631

https://tree.opentreeoflife.org/curator/study/view/ot_521?tab=trees&tree=tree1
Burleigh, J. Gordon, Rebecca T. Kimball, Edward L. Braun. 2015. Building the avian tree of life using a large-scale, sparse supermatrix. Molecular Phylogenetics and Evolution 84: 53-63
http://dx.doi.org/10.1016/j.ympev.2014.12.003

https://tree.opentreeoflife.org/curator/study/view/ot_1091?tab=trees&tree=tree1
Millicent D. Sanciangco, Kent E. Carpenter, Ricardo Betancur-R., 2016, 'Phylogenetic placement of enigmatic percomorph families (Teleostei: Percomorphaceae)', Molecular Phylogenetics and Evolution, vol. 94, pp. 565-576
http://dx.doi.org/10.1016/j.ympev.2015.10.006

https://tree.opentreeoflife.org/curator/study/view/pg_1428?tab=trees&tree=tree2855
Meredith, R.W., Janecka J., Gatesy J., Ryder O.A., Fisher C., Teeling E., Goodbla A., Eizirik E., Simao T., Stadler T., Rabosky D., Honeycutt R., Flynn J., Ingram C., Steiner C., Williams T., Robinson T., Herrick A., Westerman M., Ayoub N., Springer M., & Murphy W. 2011. Impacts of the Cretaceous Terrestrial Revolution and KPg Extinction on Mammal Diversification. Science 334 (6055): 521-524.
http://dx.doi.org/10.1126/science.1211028

https://tree.opentreeoflife.org/curator/study/view/pg_2460?tab=trees&tree=tree5285
Pyron, Robert Alexander, Frank T Burbrink, John J Wiens. 2013. A phylogeny and revised classification of Squamata, including 4161 species of lizards and snakes. BMC Evolutionary Biology 13 (1): 93.
http://dx.doi.org/10.1186/1471-2148-13-93

https://tree.opentreeoflife.org/curator/study/view/ot_1041?tab=trees&tree=tree1
Yuchi Zheng, John J. Wiens, 2016, 'Combining phylogenomic and supermatrix approaches, and a time-calibrated phylogeny for squamate reptiles (lizards and snakes) based on 52 genes and 4162 species', Molecular Phylogenetics and Evolution, vol. 94, pp. 537-547
http://dx.doi.org/10.1016/j.ympev.2015.10.009

https://tree.opentreeoflife.org/curator/study/view/ot_307?tab=trees&tree=tree1
Wright, April M., Kathleen M. Lyons, Matthew C. Brandley, David M. Hillis. 2015. Which came first: The lizard or the egg? Robustness in phylogenetic reconstruction of ancestral states. Journal of Experimental Zoology Part B: Molecular and Developmental Evolution 324 (6): 504-516
http://dx.doi.org/10.1002/jez.b.22642

https://tree.opentreeoflife.org/curator/study/view/ot_1086?tab=trees&tree=tree1
Bazinet, A. L., Mitter, K. T., Davis, D. R., Van Nieukerken, E. J., Cummings, M. P., & Mitter, C. (2017). Phylotranscriptomics resolves ancient divergences in the Lepidoptera. Systematic Entomology, 42(2), 305-316
http://dx.doi.org/10.1111/syen.12217

https://tree.opentreeoflife.org/curator/study/view/pg_2657?tab=trees&tree=tree6191
Near, Thomas J., Michael Sandel, Kristen L. Kuhn, Peter J. Unmack, Peter C. Wainwright, Wm. Leo Smith. 2012. Nuclear gene-inferred phylogenies resolve the relationships of the enigmatic Pygmy Sunfishes, Elassoma (Teleostei: Percomorpha). Molecular Phylogenetics and Evolution 63 (2): 388-395.
http://dx.doi.org/10.1016/j.ympev.2012.01.011

https://tree.opentreeoflife.org/curator/study/view/pg_1338?tab=trees&tree=tree2666
Simon, S., Strauss S., Von haeseler A., & Hadrys H. 2009. A phylogenomic approach to resolve the basal pterygote divergence. Molecular Biology and Evolution 26 (12): 2719-2730.
http://dx.doi.org/10.1093/molbev/msp191

https://tree.opentreeoflife.org/curator/study/view/pg_2813?tab=trees&tree=tree6550
Rota-Stabelli, O., L. Campbell, H. Brinkmann, G. D. Edgecombe, S. J. Longhorn, K. J. Peterson, D. Pisani, H. Philippe, M. J. Telford. 2011. A congruent solution to arthropod phylogeny: phylogenomics, microRNAs and morphology support monophyletic Mandibulata. Proceedings of the Royal Society B: Biological Sciences 278 (1703): 298-306.
http://dx.doi.org/10.1098/rspb.2010.0590

https://tree.opentreeoflife.org/curator/study/view/pg_1217?tab=trees&tree=tree2455
Delsuc, Frédéric, Georgia Tsagkogeorga, Nicolas Lartillot, andHervé Philippe. 2008. Additional molecular
support for the new chordate phylogeny. Genesis 46 (11): 592-604.
http://dx.doi.org/10.1002/dvg.20450

https://tree.opentreeoflife.org/curator/study/view/pg_2629?tab=trees&tree=tree6162
Letsch, H. and S. Simon. 2013. Insect phylogenomics: new insights on the relationships of lower neopteran orders (Polyneoptera). Systematic Entomology 38 (4): 783-793.
http://dx.doi.org/10.1111/syen.12028

https://tree.opentreeoflife.org/curator/study/view/ot_185?tab=trees&tree=tree1
Kawahara, A. Y., J. W. Breinholt. 2014. Phylogenomics provides strong evidence for relationships of butterflies and moths. Proceedings of the Royal Society B: Biological Sciences 281 (1788): 20140970-20140970.
http://dx.doi.org/10.1098/rspb.2014.0970

https://tree.opentreeoflife.org/curator/study/view/ot_1046?tab=trees&tree=tree1
Seong-il Eyun, 2017, 'Phylogenomic analysis of Copepoda (Arthropoda, Crustacea) reveals unexpected similarities with earlier proposed morphological phylogenies', BMC Evolutionary Biology, vol. 17, no. 1
http://dx.doi.org/10.1186/s12862-017-0883-5

https://tree.opentreeoflife.org/curator/study/view/pg_1337?tab=trees&tree=tree6167
Wiegmann, B., Trautwein M., Kim J., Bertone M., Winterton S., Cassel B., & Yeates D. 2009. Nuclear genes resolve the phylogeny of the holometabolous insects. BMC Biology 7(34): 34.
http://dx.doi.org/10.1186/1741-7007-7-34

https://tree.opentreeoflife.org/curator/study/view/ot_1862?tab=trees&tree=tree2
Marianne Espeland, Jesse Breinholt, Keith R. Willmott, Andrew D. Warren, Roger Vila, Emmanuel F.A. Toussaint, Sarah C. Maunsell, Kwaku Aduse-Poku, Gerard Talavera, Rod Eastwood, Marta A. Jarzyna, Robert Guralnick, David J. Lohman, Naomi E. Pierce, Akito Y. Kawahara, 2018, 'A Comprehensive and Dated Phylogenomic Analysis of Butterflies', Current Biology, vol. 28, no. 5, pp. 770-778.e5
http://dx.doi.org/10.1016/j.cub.2018.01.061

https://tree.opentreeoflife.org/curator/study/view/ot_156?tab=trees&tree=tree1
Moyle, Robert G. 2004. Phylogenetics of barbets (Aves: Piciformes) based on nuclear and mitochondrial DNA sequence data. Molecular Phylogenetics and Evolution 30 (1): 187-200.
http://dx.doi.org/10.1016/s1055-7903(03)00179-9

https://tree.opentreeoflife.org/curator/study/view/ot_1085?tab=trees&tree=tree1
Adam L. Bazinet, Michael P. Cummings, Kim T. Mitter, Charles W. Mitter, 2013, 'Can RNA-Seq Resolve the Rapid Radiation of Advanced Moths and Butterflies (Hexapoda: Lepidoptera: Apoditrysia)? An Exploratory Study', PLoS ONE, vol. 8, no. 12, p. e82615
http://dx.doi.org/10.1371/journal.pone.0082615

https://tree.opentreeoflife.org/curator/study/view/ot_1117?tab=trees&tree=tree2
Harald Letsch, Brigitte Gottsberger, Jessica L. Ware, 2016, 'Not going with the flow: a comprehensive time-calibrated phylogeny of dragonflies (Anisoptera: Odonata: Insecta) provides evidence for the role of lentic habitats on diversification', Molecular Ecology, vol. 25, no. 6, pp. 1340-1353
http://dx.doi.org/10.1111/mec.13562

https://tree.opentreeoflife.org/curator/study/view/ot_794?tab=trees&tree=tree2
Fuchs, Jérôme, Jan I. Ohlson, Per G. P. Ericson, Eric Pasquet. 2007. Synchronous intercontinental splits between assemblages of woodpeckers suggested by molecular data. Zoologica Scripta 36 (1): 11-25
http://dx.doi.org/10.1111/j.1463-6409.2006.00267.x

https://tree.opentreeoflife.org/curator/study/view/ot_1104?tab=trees&tree=tree3
Jessica Ware, Michael May, Karl Kjer, 2007, 'Phylogeny of the higher Libelluloidea (Anisoptera: Odonata): An exploration of the most speciose superfamily of dragonflies', Molecular Phylogenetics and Evolution, vol. 45, no. 1, pp. 289-310
http://dx.doi.org/10.1016/j.ympev.2007.05.027

https://tree.opentreeoflife.org/curator/study/view/pg_2913?tab=trees&tree=tree6741
Price, Trevor D., Daniel M. Hooper, Caitlyn D. Buchanan, Ulf S. Johansson, D. Thomas Tietze, Per Alström, Urban Olsson, Mousumi Ghosh-Harihar, Farah Ishtiaq, Sandeep K. Gupta, Jochen Martens, Bettina Harr, Pratap Singh, Dhananjai Mohan. 2014. Niche filling slows the diversification of Himalayan songbirds. Nature 509: 222-225.
http://dx.doi.org/10.1038/nature13272

https://tree.opentreeoflife.org/curator/study/view/pg_2688?tab=trees&tree=tree6240
Fabre, Pierre-Henri, Lionel Hautier, Dimitar Dimitrov, Emmanuel J P Douzery. 2012. A glimpse on the pattern of rodent diversification: a phylogenetic approach. BMC Evolutionary Biology 12 (1): 88.
http://dx.doi.org/10.1186/1471-2148-12-88

https://tree.opentreeoflife.org/curator/study/view/ot_1641?tab=trees&tree=tree1
Michael E. Alfaro, Brant C. Faircloth, Richard C. Harrington, Laurie Sorenson, Matt Friedman, Christine E. Thacker, Carl H. Oliveros, David Černý, Thomas J. Near, 2018, 'Explosive diversification of marine fishes at the Cretaceous–Palaeogene boundary', Nature Ecology & Evolution, vol. 2, no. 4, pp. 688-696
http://dx.doi.org/10.1038/s41559-018-0494-6

https://tree.opentreeoflife.org/curator/study/view/pg_2853?tab=trees&tree=tree6624
McGuire, Jimmy A., Christopher C. Witt, J.V. Remsen, Ammon Corl, Daniel L. Rabosky, Douglas L. Altshuler, Robert Dudley. 2014. Molecular phylogenetics and the diversification of hummingbirds. Current Biology 24 (8): 910-916.
http://dx.doi.org/10.1016/j.cub.2014.03.016

https://tree.opentreeoflife.org/curator/study/view/pg_2822?tab=trees&tree=tree6569
Parfrey, L. W., D. J. G. Lahr, A. H. Knoll, L. A. Katz. 2011. Estimating the timing of early eukaryotic diversification with multigene molecular clocks. Proceedings of the National Academy of Sciences 108 (33): 13624-13629.
http://dx.doi.org/10.1073/pnas.1110633108

https://tree.opentreeoflife.org/curator/study/view/ot_1366?tab=trees&tree=Tr98763
Mark S. Springer, Christopher A. Emerling, Robert W. Meredith, Jan E. Janečka, Eduardo Eizirik, William J. Murphy, 2017, 'Waking the undead: Implications of a soft explosive model for the timing of placental mammal diversification', Molecular Phylogenetics and Evolution, vol. 106, pp. 86-102
http://dx.doi.org/10.1016/j.ympev.2016.09.017

https://tree.opentreeoflife.org/curator/study/view/pg_2651?tab=trees&tree=tree6177
Betancur-R., Ricardo, Richard E. Broughton, Edward O. Wiley, Kent Carpenter, J. Andrés López, Chenhong Li, Nancy I. Holcroft, Dahiana Arcila, Millicent Sanciangco, James C Cureton II, Feifei Zhang, Thaddaeus Buser, Matthew A. Campbell, Jesus A Ballesteros, Adela Roa-Varon, Stuart Willis, W. Calvin Borden, Thaine Rowley, Paulette C. Reneau, Daniel J. Hough, Guoqing Lu, Terry Grande, Gloria Arratia, Guillermo Ortí. 2013. The Tree of Life and a New Classification of Bony Fishes. PLoS Currents
http://dx.doi.org/10.1371/currents.tol.53ba26640df0ccaee75bb165c8c26288

https://tree.opentreeoflife.org/curator/study/view/ot_1286?tab=trees&tree=tree1
Brian Tilston Smith, Robert W. Bryson, William M. Mauck, Jaime Chaves, Mark B. Robbins, Alexandre Aleixo, John Klicka, 2018, 'Species delimitation and biogeography of the gnatcatchers and gnatwrens (Aves: Polioptilidae)', Molecular Phylogenetics and Evolution, vol. 126, pp. 45-57
http://dx.doi.org/10.1016/j.ympev.2018.03.012

https://tree.opentreeoflife.org/curator/study/view/pg_2604?tab=trees&tree=tree6043
Hedtke, S.M., Patiny S., & Danforth B. 2013. The bee tree of life: a supermatrix approach to apoid phylogeny and biogeography. BMC Evolutionary Biology 13: 138.
http://dx.doi.org/10.1186/1471-2148-13-138

https://tree.opentreeoflife.org/curator/study/view/ot_504?tab=trees&tree=tree3
Dufort, Matthew J. 2015. An augmented supermatrix phylogeny of the avian family Picidae reveals uncertainty deep in the family tree. Molecular Phylogenetics and Evolution
http://dx.doi.org/10.1016/j.ympev.2015.08.025

https://tree.opentreeoflife.org/curator/study/view/ot_508?tab=trees&tree=tree2
dos Reis, Mario, Yuttapong Thawornwattana, Konstantinos Angelis, Maximilian J. Telford, Philip C.J. Donoghue, Ziheng Yang. 2015. Uncertainty in the timing of origin of animals and the limits of precision in molecular timescales. Current Biology
http://dx.doi.org/10.1016/j.cub.2015.09.066

https://tree.opentreeoflife.org/curator/study/view/pg_2576?tab=trees&tree=tree5975
Near, T. J., A. Dornburg, R. I. Eytan, B. P. Keck, W. L. Smith, K. L. Kuhn, J. A. Moore, S. A. Price, F. T. Burbrink, M. Friedman, P. C. Wainwright, 2013, 'Phylogeny and tempo of diversification in the superradiation of spiny-rayed fishes', Proceedings of the National Academy of Sciences, vol. 110, no. 31, pp. 12738-12743
http://dx.doi.org/10.1073/pnas.1304661110

https://tree.opentreeoflife.org/curator/study/view/ot_520?tab=trees&tree=tree1
Jønsson, Knud Andreas, Pierre-Henri Fabre, Jonathan D. Kennedy, Ben G. Holt, Michael K. Borregaard, Carsten Rahbek, Jon Fjeldså. 2016. A supermatrix phylogeny of corvoid passerine birds (Aves: Corvides). Molecular Phylogenetics and Evolution 94: 87-94.
http://dx.doi.org/10.1016/j.ympev.2015.08.020

https://tree.opentreeoflife.org/curator/study/view/ot_1512?tab=trees&tree=Tr55545
Brusatte S.L., Benton M.J., Desojo J.B., & Langer M.C. 2010. The higher-level phylogeny of Archosauria (Tetrapoda: Diapsida). Journal of Systematic Palaeontology, 8(1): 3-47.
http://dx.doi.org/10.1080/14772010903537732

https://tree.opentreeoflife.org/curator/study/view/pg_2710?tab=trees&tree=tree6291
Ryan, J. F., K. Pang, C. E. Schnitzler, A.-D. Nguyen, R. T. Moreland, D. K. Simmons, B. J. Koch, W. R. Francis, P. Havlak, S. A. Smith, N. H. Putnam, S. H. D. Haddock, C. W. Dunn, T. G. Wolfsberg, J. C. Mullikin, M. Q. Martindale, A. D. Baxevanis. 2013. The genome of the ctenophore Mnemiopsis leidyi and its implications for cell type evolution. Science 342 (6164): 1242592-1242592.
http://dx.doi.org/10.1126/science.1242592

https://tree.opentreeoflife.org/curator/study/view/pg_2551?tab=trees&tree=tree6180
Wainwright, P.C., Smith W.L., Price S.A., Tang K.L., Sparks J.S., Ferry L.A., Kuhn K.L., Eytan R.I., & Near T.J. 2012. The evolution of pharyngognathy: a phylogenetic and functional appraisal of the pharyngeal jaw key innovation in labroid fishes and beyond. Systematic Biology 61 (6): 1001-1027.
http://dx.doi.org/10.1093/sysbio/sys060

https://tree.opentreeoflife.org/curator/study/view/pg_2804?tab=trees&tree=tree6511
Kimball, Rebecca T., Colette M. St. Mary, Edward L. Braun. 2011. A macroevolutionary perspective on multiple sexual traits in the Phasianidae (Galliformes). International Journal of Evolutionary Biology 2011: 1-16.
http://dx.doi.org/10.4061/2011/423938

https://tree.opentreeoflife.org/curator/study/view/pg_2814?tab=trees&tree=tree6551
Rota-Stabelli, O., Lartillot, N., Philippe, H., and Pisani, D. 2013. Serine codon-usage bias in deep phylogenomics: Pancrustacean relationships as a case study. Systematic Biology 62 (1): 121-133.
http://dx.doi.org/10.1093/sysbio/sys077

https://tree.opentreeoflife.org/curator/study/view/ot_1047?tab=trees&tree=tree1
T. H. Oakley, J. M. Wolfe, A. R. Lindgren, A. K. Zaharoff, 2012, 'Phylotranscriptomics to Bring the Understudied into the Fold: Monophyletic Ostracoda, Fossil Placement, and Pancrustacean Phylogeny', Molecular Biology and Evolution, vol. 30, no. 1, pp. 215-233
http://dx.doi.org/10.1093/molbev/mss216

https://tree.opentreeoflife.org/curator/study/view/ot_848?tab=trees&tree=tree1
Irby J. Lovette, Brynn V. McCleery, Amanda L. Talaba, Dustin R. Rubenstein, 2008, 'A complete species-level molecular phylogeny for the “Eurasian” starlings (Sturnidae: Sturnus, Acridotheres, and allies): Recent diversification in a highly social and dispersive avian group', Molecular Phylogenetics and Evolution, vol. 47, no. 1, pp. 251-260
http://dx.doi.org/10.1016/j.ympev.2008.01.020

https://tree.opentreeoflife.org/curator/study/view/pg_2591?tab=trees&tree=tree6024
Lovette, Irby J., Jorge L. Pérez-Emán, John P. Sullivan, Richard C. Banks, Isabella Fiorentino, Sergio Córdoba-Córdoba, María Echeverry-Galvis, F. Keith Barker, Kevin J. Burns, John Klicka, Scott M. Lanyon, Eldredge Bermingham. 2010. A comprehensive multilocus phylogeny for the wood-warblers and a revised classification of the Parulidae (Aves). Molecular Phylogenetics and Evolution 57 (2): 753-770.
http://dx.doi.org/10.1016/j.ympev.2010.07.018

https://tree.opentreeoflife.org/curator/study/view/ot_1431?tab=trees&tree=Tr55921
Dongyu H., Lianhai H., Lijun Z., & Xing X. 2010. A pre-Archaeopteryx troodontid theropod from China with long feathers on the metatarsus. Nature, 461: 640-643.
http://dx.doi.org/10.1038/nature08322

https://tree.opentreeoflife.org/curator/study/view/ot_1513?tab=trees&tree=Tr55903
Zanno L.E. 2010. A taxonomic and phylogenetic re-evaluation of Therizinosauria (Dinosauria: Maniraptora). Journal of Systematic Palaeontology, 8(4): 37-41.
http://dx.doi.org/10.1080/14772019.2010.488045

https://tree.opentreeoflife.org/curator/study/view/ot_1510?tab=trees&tree=Tr55267
Carrano M.T., Benson R.B., & Sampson S.D. 2012. The phylogeny of Tetanurae (Dinosauria: Theropoda). Journal of Systematic Palaeontology, 10(2): 211-300.
http://dx.doi.org/10.1080/14772019.2011.630927

https://tree.opentreeoflife.org/curator/study/view/ot_1517?tab=trees&tree=Tr60095
Andres B.B., Clark J.M., & Xu X. 2012. A new rhamphorhynchid pterosaur from the Upper Jurassic of Xinjiang, China, and the phylogenetic relationships of basal pterosaurs. Journal of Vertebrate Paleontology, 30(1): 163-187.
http://dx.doi.org/10.1080/02724630903409220

https://tree.opentreeoflife.org/curator/study/view/ot_278?tab=trees&tree=tree1
Tedford, Richard H.; Wang, Xiaoming; Taylor, Beryl E. (2009). Phylogenetic systematics of the North American fossil Caninae (Carnivora, Canidae). Bulletin of the American Museum of Natural History, no. 325. http://hdl.handle.net/2246/5999

Wang, Xiaoming; Tedford, Richard H.; Taylor, Beryl E. (1999). Phylogenetic systematics of the Borophaginae (Carnivora, Canidae). Bulletin of the American Museum of Natural History, no. 243. http://hdl.handle.net/2246/1588

Wang, Xiaoming (1994). Phylogenetic systematics of the Hesperocyoninae (Carnivora, Canidae). Bulletin of the  American Museum of Natural History, no. 221. http://hdl.handle.net/2246/829

http://hdl.handle.net/2246/5999

https://tree.opentreeoflife.org/curator/study/view/pg_2845?tab=trees&tree=tree6606
Pasquet,  Eric, F. Keith Barker, Jochen Martens, Annie Tillier, Corinne Cruaud, Alice Cibois. 2014.  Evolution within the nuthatches (Sittidae: Aves, Passeriformes): molecular phylogeny, biogeography, and ecological perspectives. Journal of Ornithology 155 (3):755-765.
http://dx.doi.org/10.1007/s10336-014-1063-7

https://tree.opentreeoflife.org/curator/study/view/ot_126?tab=trees&tree=tree1
Krajewski, Carey, Justin T. Sipiorski, Frank E. Anderson. 2010. Complete mitochondrial genome sequences and the phylogeny of cranes (Gruiformes: Gruidae). The Auk 127 (2): 440-452.
http://dx.doi.org/10.1525/auk.2009.09045

https://tree.opentreeoflife.org/curator/study/view/ot_171?tab=trees&tree=tree1
Spellman, GM, A Cibois, RG Moyle, K Winker, F Keith Barker. 2008. Clarifying the systematics of an enigmatic avian lineage: What is a bombycillid?. Molecular Phylogenetics and Evolution 49 (3): 1036-1040.
http://dx.doi.org/10.1016/j.ympev.2008.09.006

https://tree.opentreeoflife.org/curator/study/view/pg_1764?tab=trees&tree=tree6299
Kennedy, M.,  S.A. Taylor, P. Nádvorník, H. Spencer. 2013. The phylogenetic relationships of the extant pelicans inferred from DNA sequence data. Molecular Phylogenetics and Evolution 66 (1): 215-222.
http://dx.doi.org/10.1016/j.ympev.2012.09.034

https://tree.opentreeoflife.org/curator/study/view/ot_812?tab=trees&tree=tree4
Johnson, Jeff A., Joseph W. Brown, Jérôme Fuchs, David P. Mindell, 2016, 'Multi-locus phylogenetic inference among New World Vultures (Aves: Cathartidae)', Molecular Phylogenetics and Evolution, vol. 105, pp. 193-199
http://dx.doi.org/10.1016/j.ympev.2016.08.025

https://tree.opentreeoflife.org/curator/study/view/ot_864?tab=trees&tree=tree1
Ogawa, Lisa M., Paulo C. Pulgarin, Donald A. Vance, Jon Fjeldså, Marcel van Tuinen. 2015. Opposing demographic histories reveal rapid evolution in grebes (Aves: Podicipedidae). The Auk 132 (4): 771-786
http://dx.doi.org/10.1642/auk-14-259.1

https://tree.opentreeoflife.org/curator/study/view/ot_177?tab=trees&tree=tree1
Dor, Roi, Rebecca J. Safran, Frederick H. Sheldon, David W. Winkler, Irby J. Lovette. 2010. Phylogeny of the genus Hirundo and the Barn Swallow subspecies complex. Molecular Phylogenetics and Evolution 56 (1): 409-418.
http://dx.doi.org/10.1016/j.ympev.2010.02.008

https://tree.opentreeoflife.org/curator/study/view/ot_172?tab=trees&tree=tree3
Päckert, Martin, Jochen Martens, Yue-Hua Sun, Lucia Liu Severinghaus, Alexander A. Nazarenko, Ji Ting, Till Töpfer, Dieter Thomas Tietze. 2011. Horizontal and elevational phylogeographic patterns of Himalayan and Southeast Asian forest passerines (Aves: Passeriformes). Journal of Biogeography 39 (3): 556-573.
http://dx.doi.org/10.1111/j.1365-2699.2011.02606.x

https://tree.opentreeoflife.org/curator/study/view/ot_104?tab=trees&tree=tree1
Boertmann, D. 1990. Phylogeny of the divers, family Gaviidae (Aves). Steenstrupia 16: 21-36.


https://tree.opentreeoflife.org/curator/study/view/ot_1592?tab=trees&tree=tree1
Daniel L. Rabosky, Jonathan Chang, Pascal O. Title, Peter F. Cowman, Lauren Sallan, Matt Friedman, Kristin Kaschner, Cristina Garilao, Thomas J. Near, Marta Coll, Michael E. Alfaro, 2018, 'An inverse latitudinal gradient in speciation rate for marine fishes', Nature, vol. 559, no. 7714, pp. 392-395
http://dx.doi.org/10.1038/s41586-018-0273-1

https://tree.opentreeoflife.org/curator/study/view/ot_1164?tab=trees&tree=tree2
Yan-Jie Feng, David C. Blackburn, Dan Liang, David M. Hillis, David B. Wake, David C. Cannatella, Peng Zhang, 2017, 'Phylogenomics reveals rapid, simultaneous diversification of three major clades of Gondwanan frogs at the Cretaceous–Paleogene boundary', Proceedings of the National Academy of Sciences, p. 201704632
http://dx.doi.org/10.1073/pnas.1704632114

https://tree.opentreeoflife.org/curator/study/view/ot_1164?tab=trees&tree=tree3
Yan-Jie Feng, David C. Blackburn, Dan Liang, David M. Hillis, David B. Wake, David C. Cannatella, Peng Zhang, 2017, 'Phylogenomics reveals rapid, simultaneous diversification of three major clades of Gondwanan frogs at the Cretaceous–Paleogene boundary', Proceedings of the National Academy of Sciences, p. 201704632
http://dx.doi.org/10.1073/pnas.1704632114

https://tree.opentreeoflife.org/curator/study/view/pg_423?tab=trees&tree=tree2857
Pyron, R.A., & Wiens J.J. 2011. A large-scale phylogeny of Amphibia including over 2800 species, and a revised classification of extant frogs, salamanders, and caecilians. Molecular Phylogenetics Evolution 61 (2): 543-583.
http://dx.doi.org/10.1016/j.ympev.2011.06.012

https://tree.opentreeoflife.org/curator/study/view/ot_505?tab=trees&tree=tree1
Tod W. Reeder, Ted M. Townsend, Daniel G. Mulcahy, Brice P. Noonan, Perry L. Wood, Jack W. Sites, John J. Wiens, 2015, 'Integrated Analyses Resolve Conflicts over Squamate Reptile Phylogeny and Reveal Unexpected Placements for Fossil Taxa', PLOS ONE, vol. 10, no. 3, p. e0118199
http://dx.doi.org/10.1371/journal.pone.0118199

https://tree.opentreeoflife.org/curator/study/view/ot_290?tab=trees&tree=tree1
Selvatti, Alexandre Pedro, Luiz Pedreira Gonzaga, Claudia Augusta de Moraes Russo. 2015. A Paleogene origin for crown passerines and the diversification of the Oscines in the New World. Molecular Phylogenetics and Evolution 88: 1-15.
http://dx.doi.org/10.1016/j.ympev.2015.03.018

https://tree.opentreeoflife.org/curator/study/view/ot_783?tab=trees&tree=tree1
Moyle, Robert G., Carl H. Oliveros, Michael J. Andersen, Peter A. Hosner, Brett W. Benz, Joseph D. Manthey, Scott L. Travers, Rafe M. Brown, Brant C. Faircloth. 2016. Tectonic collision and uplift of Wallacea triggered the global songbird radiation. Nature Communications 7: 12709
http://dx.doi.org/10.1038/ncomms12709

https://tree.opentreeoflife.org/curator/study/view/pg_2015?tab=trees&tree=tree4152
Odeen, A., Hastad O., & Alstrom P. 2011. Evolution of ultraviolet vision in the largest avian radiation - the passerines. BMC Evolutionary Biology 11 (313): 1-8.
http://dx.doi.org/10.1186/1471-2148-11-313

https://tree.opentreeoflife.org/curator/study/view/ot_531?tab=trees&tree=tree1
Claramunt, Santiago, Joel Cracraft. 2015. A new time tree reveals Earth historys imprint on the evolution of modern birds. Science Advances 1 (11): e1501005-e1501005
http://dx.doi.org/10.1126/sciadv.1501005

https://tree.opentreeoflife.org/curator/study/view/ot_782?tab=trees&tree=Tr48736
Beresford, P., F.K. Barker, P.G. Ryan, T.M. Crowe. 2005. African endemics span the tree of songbirds (Passeri): molecular systematics of several evolutionary 'enigmas'. Proceedings of the Royal Society B: Biological Sciences 272 (1565): 849-858
http://dx.doi.org/10.1098/rspb.2004.2997

https://tree.opentreeoflife.org/curator/study/view/ot_1844?tab=trees&tree=tree1
Martin Stervander, Peter G. Ryan, Martim Melo, Bengt Hansson, 2019, 'The origin of the world’s smallest flightless bird, the Inaccessible Island Rail Atlantisia rogersi (Aves: Rallidae)', Molecular Phylogenetics and Evolution, vol. 130, pp. 92-98
http://dx.doi.org/10.1016/j.ympev.2018.10.007

https://tree.opentreeoflife.org/curator/study/view/pg_2575?tab=trees&tree=tree5974
Barker, F. K., K. J. Burns, J. Klicka, S. M. Lanyon, I. J. Lovette. 2013. Going to extremes: contrasting rates of diversification in a recent radiation of New World passerine birds. Systematic Biology 62 (2): 298-320.
http://dx.doi.org/10.1093/sysbio/sys094

https://tree.opentreeoflife.org/curator/study/view/ot_770?tab=trees&tree=tree1
Barker, F. Keith, Kevin J. Burns, John Klicka, Scott M. Lanyon, Irby J. Lovette. 2015. New insights into New World biogeography: An integrated view from the phylogeny of blackbirds, cardinals, sparrows, tanagers, warblers, and allies. The Auk 132 (2): 333-348.
http://dx.doi.org/10.1642/auk-14-110.1

https://tree.opentreeoflife.org/curator/study/view/pg_2924?tab=trees&tree=tree6755
Klicka, John, F. Keith Barker, Kevin J. Burns, Scott M. Lanyon, Irby J. Lovette, Jaime A. Chaves, Robert W. Bryson. 2014. A comprehensive multilocus assessment of sparrow (Aves: Passerellidae) relationships. Molecular Phylogenetics and Evolution 77: 177–182.
http://dx.doi.org/10.1016/j.ympev.2014.04.025

https://tree.opentreeoflife.org/curator/study/view/ot_841?tab=trees&tree=tree1
Simone Treplin, Romy Siegert, Christoph Bleidorn, Hazell Shokellu Thompson, Roger Fotso, Ralph Tiedemann, 2008, 'Molecular phylogeny of songbirds (Aves: Passeriformes) and the relative utility of common nuclear marker loci', Cladistics, vol. 24, no. 3, pp. 328-349
http://dx.doi.org/10.1111/j.1096-0031.2007.00178.x

https://tree.opentreeoflife.org/curator/study/view/pg_2913?tab=trees&tree=tree6742
Price, Trevor D., Daniel M. Hooper, Caitlyn D. Buchanan, Ulf S. Johansson, D. Thomas Tietze, Per Alström, Urban Olsson, Mousumi Ghosh-Harihar, Farah Ishtiaq, Sandeep K. Gupta, Jochen Martens, Bettina Harr, Pratap Singh, Dhananjai Mohan. 2014. Niche filling slows the diversification of Himalayan songbirds. Nature 509: 222-225.
http://dx.doi.org/10.1038/nature13272

https://tree.opentreeoflife.org/curator/study/view/ot_534?tab=trees&tree=tree1
Ksepka, Daniel T., Matthew J. Phillips. 2015. Avian diversification patterns across the K-Pg boundary: influence of calibrations, datasets, and model misspecification. Annals of the Missouri Botanical Garden 100 (4): 300-328
http://dx.doi.org/10.3417/2014032

https://tree.opentreeoflife.org/curator/study/view/ot_757?tab=trees&tree=Tr70438
Ericson, Per GP, Seraina Klopfstein, Martin Irestedt, Jacqueline MT Nguyen and Johan AA Nylander. 2014. Dating the diversification of the major lineages of Passeriformes (Aves). BMC Evolutionary Biology, 14(8).
http://dx.doi.org/10.1186/1471-2148-14-8

https://tree.opentreeoflife.org/curator/study/view/pg_420?tab=trees&tree=tree522
Hackett, S. J., R. T. Kimball, S. Reddy, R. C. K. Bowie, E. L. Braun, M. J. Braun, J. L. Chojnowski, W. A. Cox, K.-L. Han, J. Harshman, C. J. Huddleston, B. D. Marks, K. J. Miglia, W. S. Moore, F. H. Sheldon, D. W. Steadman, C. C. Witt, T. Yuri. 2008. A Phylogenomic Study of Birds Reveals Their Evolutionary History. Science 320 (5884): 1763-1768.
http://dx.doi.org/10.1126/science.1157704

https://tree.opentreeoflife.org/curator/study/view/ot_412?tab=trees&tree=Tr77157
Barker, F.K. 2014. Mitogenomic data resolve basal relationships among passeriform and passeridan birds. Molecular Phylogenetics and Evolution 79: 313-324.
http://dx.doi.org/10.1016/j.ympev.2014.06.011

https://tree.opentreeoflife.org/curator/study/view/ot_500?tab=trees&tree=tree1
Prum, Richard O., Jacob S. Berv, Alex Dornburg, Daniel J. Field, Jeffrey P. Townsend, Emily Moriarty Lemmon, Alan R. Lemmon. 2015. A comprehensive phylogeny of birds (Aves) using targeted next-generation DNA sequencing. Nature 526, (7574): 569-573
http://dx.doi.org/10.1038/nature15697

https://tree.opentreeoflife.org/curator/study/view/ot_532?tab=trees&tree=tree1
Gibb, Gillian C., Ryan England, Gerrit Hartig, P.A. (Trish) McLenachan, Briar L. Taylor Smith, Bennet J. McComish, Alan Cooper, David Penny. 2015. New Zealand passerines help clarify the diversification of major songbird lineages during the Oligocene. Genome Biology and Evolution 7 (11): 2983-2995.
http://dx.doi.org/10.1093/gbe/evv196

https://tree.opentreeoflife.org/curator/study/view/pg_2404?tab=trees&tree=tree5068
Barker, F. Keith, Alice Cibois, Peter Schikler, Julie Feinstein, Joel Cracraft. 2004. Phylogeny and diversification of the largest avian radiation. Proceedings of the National Academy of Sciences of the United States of America 101 (30): 11040-11045.
http://dx.doi.org/10.1073/pnas.0401892101

https://tree.opentreeoflife.org/curator/study/view/pg_2404?tab=trees&tree=tree6388
Barker, F. Keith, Alice Cibois, Peter Schikler, Julie Feinstein, Joel Cracraft. 2004. Phylogeny and diversification of the largest avian radiation. Proceedings of the National Academy of Sciences of the United States of America 101 (30): 11040-11045.
http://dx.doi.org/10.1073/pnas.0401892101

https://tree.opentreeoflife.org/curator/study/view/ot_1053?tab=trees&tree=tree1
Ralph S. Peters, Lars Krogmann, Christoph Mayer, Alexander Donath, Simon Gunkel, Karen Meusemann, Alexey Kozlov, Lars Podsiadlowski, Malte Petersen, Robert Lanfear, Patricia A. Diez, John Heraty, Karl M. Kjer, Seraina Klopfstein, Rudolf Meier, Carlo Polidori, Thomas Schmitt, Shanlin Liu, Xin Zhou, Torsten Wappler, Jes Rust, Bernhard Misof, Oliver Niehuis, 2017, 'Evolutionary History of the Hymenoptera', Current Biology
http://dx.doi.org/10.1016/j.cub.2017.01.027

https://tree.opentreeoflife.org/curator/study/view/ot_1059?tab=trees&tree=tree1
Michael G. Branstetter, Bryan N. Danforth, James P. Pitts, Brant C. Faircloth, Philip S. Ward, Matthew L. Buffington, Michael W. Gates, Robert R. Kula, Seán G. Brady, 2017, 'Phylogenomic Insights into the Evolution of Stinging Wasps and the Origins of Ants and Bees', Current Biology, vol. 27, no. 7, pp. 1019-1025
http://dx.doi.org/10.1016/j.cub.2017.03.027

https://tree.opentreeoflife.org/curator/study/view/pg_2692?tab=trees&tree=tree6245
Zuccon, Dario, Robert Prŷs-Jones, Pamela C. Rasmussen, and Per G.P. Ericson. 2012. The phylogenetic relationships and generic limits of finches (Fringillidae). Molecular Phylogenetics and Evolution 62 (2): 581-596.
http://dx.doi.org/10.1016/j.ympev.2011.10.002

https://tree.opentreeoflife.org/curator/study/view/ot_854?tab=trees&tree=tree1
Ulf S. Johansson, Jon Fjeldså, Rauri C.K. Bowie, 2008, 'Phylogenetic relationships within Passerida (Aves: Passeriformes): A review and a new molecular phylogeny based on three nuclear intron markers', Molecular Phylogenetics and Evolution, vol. 48, no. 3, pp. 858-876
http://dx.doi.org/10.1016/j.ympev.2008.05.029

https://tree.opentreeoflife.org/curator/study/view/ot_111?tab=trees&tree=tree1
Alstrom, P., D. M. Hooper, Y. Liu, U. Olsson, D. Mohan, M. Gelang, H. Le Manh, J. Zhao, F. Lei, T. D. Price. 2014. Discovery of a relict lineage and monotypic family of passerine birds. Biology Letters 10 (3): 20131067-20131067.
http://dx.doi.org/10.1098/rsbl.2013.1067

https://tree.opentreeoflife.org/curator/study/view/pg_2573?tab=trees&tree=tree5959
Crawford, N. G., B. C. Faircloth, J. E. McCormack, R. T. Brumfield, K. Winker, T. C. Glenn. 2012. More than 1000 ultraconserved elements provide evidence that turtles are the sister group of archosaurs. Biology Letters 8 (5): 783-786.
http://dx.doi.org/10.1098/rsbl.2012.0331

https://tree.opentreeoflife.org/curator/study/view/ot_328?tab=trees&tree=tree1
Nyakatura, Katrin, Olaf RP Bininda-Emonds. 2012. Updating the evolutionary history of Carnivora (Mammalia): a new species-level supertree complete with divergence time estimates. BMC Biology 10 (1): 12
http://dx.doi.org/10.1186/1741-7007-10-12

https://tree.opentreeoflife.org/curator/study/view/ot_1410?tab=trees&tree=Tr97099
White N.D., Barrowclough G.F., Groth J.G., & Braun M. 2016. A Multi-Gene Estimate of Higher-Level Phylogenetic Relationships Among Nightjars (Aves: Caprimulgidae). Ornitologia Neotropical, 27: 223-236.


https://tree.opentreeoflife.org/curator/study/view/pg_2872?tab=trees&tree=tree6665
Han, Kin-Lan, Mark B. Robbins, Michael J. Braun. 2010. A multi-gene estimate of phylogeny in the nightjars and nighthawks (Caprimulgidae). Molecular Phylogenetics and Evolution 55 (2): 443-453.
http://dx.doi.org/10.1016/j.ympev.2010.01.023

https://tree.opentreeoflife.org/curator/study/view/pg_2707?tab=trees&tree=tree6281
Powell, Alexis F.L.A., F. Keith Barker, Scott M. Lanyon, Kevin J. Burns, John Klicka, Irby J. Lovette. 2014. A comprehensive species-level molecular phylogeny of the New World blackbirds (Icteridae). Molecular Phylogenetics and Evolution 71: 94-112.
http://dx.doi.org/10.1016/j.ympev.2013.11.009

https://tree.opentreeoflife.org/curator/study/view/ot_753?tab=trees&tree=tree1
Wang, Ning, Rebecca T. Kimball, Edward L. Braun, Bin Liang, Zhengwang Zhang. 2016. Ancestral range reconstruction of Galliformes: the effects of topology and taxon sampling. Journal of Biogeography
http://dx.doi.org/10.1111/jbi.12782

https://tree.opentreeoflife.org/curator/study/view/pg_2577?tab=trees&tree=tree5980
Wang, Ning, Rebecca T. Kimball, Edward L. Braun, Bin Liang, Zhengwang Zhang. 2013. Assessing phylogenetic relationships among Galliformes: a multigene phylogeny with expanded taxon sampling in Phasianidae. PLoS ONE 8 (5): e64312.
http://dx.doi.org/10.1371/journal.pone.0064312

https://tree.opentreeoflife.org/curator/study/view/ot_1002?tab=trees&tree=tree1
Hosner, Peter A. Brant C. Faircloth, Travis C. Glenn, Edward L. Braun, Rebecca T. Kimball. 2015. Avoiding missing data biases in phylogenomic inference: an empirical study in the landfowl (Aves: Galliformes). Molecular Biology and Evolution 33 (4): 1110-1125
http://dx.doi.org/10.1093/molbev/msv347

https://tree.opentreeoflife.org/curator/study/view/ot_1345?tab=trees&tree=Tr109614
Worthy T., Mitri M., Handley W.D., Lee M.S., Anderson A., & Sand C. 2016. Osteology Supports a Stem-Galliform Affinity for the Giant Extinct Flightless Bird Sylviornis neocaledoniae (Sylviornithidae, Galloanseres). PLoS ONE, 11(3): e0150871.
http://dx.doi.org/10.1371/journal.pone.0150871

https://tree.opentreeoflife.org/curator/study/view/ot_116?tab=trees&tree=tree1
Hugall, Andrew F., Devi Stuart-Fox. 2012. Accelerated speciation in colour-polymorphic birds. Nature 485 (7400): 631-634.
http://dx.doi.org/10.1038/nature11050

https://tree.opentreeoflife.org/curator/study/view/ot_867?tab=trees&tree=tree2
Jérôme Fuchs, Jean-Marc Pons, Steven M Goodman, Vincent Bretagnolle, Martim Melo, Rauri CK Bowie, David Currie, Roger Safford, Munir Z Virani, Simon Thomsett, Alawi Hija, Corinne Cruaud, Eric Pasquet, 2008, 'Tracing the colonization history of the Indian Ocean scops-owls (Strigiformes: Otus) with further insight into the spatio-temporal origin of the Malagasy avifauna', BMC Evolutionary Biology, vol. 8, no. 1, p. 197
http://dx.doi.org/10.1186/1471-2148-8-197

https://tree.opentreeoflife.org/curator/study/view/ot_773?tab=trees&tree=tree1
Wood, Jamie R., Kieren J. Mitchell, R. Paul Scofield, Vanesa L. De Pietri, Nicolas J. Rawlence, Alan Cooper. 2016. Phylogenetic relationships and terrestrial adaptations of the extinct laughing owl, Sceloglaux albifacies (Aves: Strigidae). Zoological Journal of the Linnean Society
http://dx.doi.org/10.1111/zoj.12483

https://tree.opentreeoflife.org/curator/study/view/pg_2812?tab=trees&tree=tree6545
Lartillot, Nicolas, Frédéric Delsuc. 2012. Joint reconstruction of divergence times and life-history evolution in placental mammals using a phylogenetic covariance model. Evolution 66 (6): 1773-1787.
http://dx.doi.org/10.1111/j.1558-5646.2011.01558.x

https://tree.opentreeoflife.org/curator/study/view/pg_2647?tab=trees&tree=tree6169
O'Leary, M. A., J. I. Bloch, J. J. Flynn, T. J. Gaudin, A. Giallombardo, N. P. Giannini, S. L. Goldberg, B. P. Kraatz, Z.-X. Luo, J. Meng, X. Ni, M. J. Novacek, F. A. Perini, Z. S. Randall, G. W. Rougier, E. J. Sargis, M. T. Silcox, N. B. Simmons, M. Spaulding, P. M. Velazco, M. Weksler, J. R. Wible, A. L. Cirranello. 2013. The placental mammal ancestor and the post-K-Pg radiation of placentals. Science 339 (6120): 662-667.
http://dx.doi.org/10.1126/science.1229237

https://tree.opentreeoflife.org/curator/study/view/pg_1887?tab=trees&tree=tree6629
Lerner, H., M. Klaver, D. Mindell. 2008. Molecular phylogenetics of the Buteonine birds of prey (Accipitridae). The Auk 125 (2): 304-315.
http://dx.doi.org/10.1525/auk.2008.06161

https://tree.opentreeoflife.org/curator/study/view/ot_837?tab=trees&tree=tree1
Robert G. Moyle, 2006, 'A MOLECULAR PHYLOGENY OF KINGFISHERS (ALCEDINIDAE) WITH INSIGHTS INTO EARLY BIOGEOGRAPHIC HISTORY', The Auk, vol. 123, no. 2, p. 487
http://dx.doi.org/10.1642/0004-8038(2006)123[487:ampoka]2.0.co;2

https://tree.opentreeoflife.org/curator/study/view/ot_1098?tab=trees&tree=tree1
Kawahara AY, Mignault AA, Regier JC, Kitching IJ, Mitter C (2009) 'Phylogeny and Biogeography of Hawkmoths (Lepidoptera: Sphingidae): Evidence from Five Nuclear Genes., PLoS ONE, vol. 4, no. 5, p. e5719
http://dx.doi.org/10.1371/journal.pone.0005719

https://tree.opentreeoflife.org/curator/study/view/ot_855?tab=trees&tree=tree1
Martin Irestedt, Jérôme Fuchs, Knud A. Jønsson, Jan I. Ohlson, Eric Pasquet, Per G.P. Ericson, 2008, 'The systematic affinity of the enigmatic Lamprolia victoriae (Aves: Passeriformes)—An example of avian dispersal between New Guinea and Fiji over Miocene intermittent land bridges?', Molecular Phylogenetics and Evolution, vol. 48, no. 3, pp. 1218-1222
http://dx.doi.org/10.1016/j.ympev.2008.05.038

https://tree.opentreeoflife.org/curator/study/view/pg_2702?tab=trees&tree=tree6274
Aggerbeck, Marie, Jon Fjeldså, Les Christidis, Pierre-Henri Fabre, Knud Andreas Jønsson. 2014. Resolving deep lineage divergences in core corvoid passerine birds supports a proto-Papuan island origin. Molecular Phylogenetics and Evolution 70: 272-285.
http://dx.doi.org/10.1016/j.ympev.2013.09.027

https://tree.opentreeoflife.org/curator/study/view/ot_843?tab=trees&tree=tree3
Rheindt, Frank E., Janette A. Norman, Les Christidis. 2008. Phylogenetic relationships of tyrant-flycatchers (Aves: Tyrannidae), with an emphasis on the elaeniine assemblage. Molecular Phylogenetics and Evolution 46 (1): 88-101
http://dx.doi.org/10.1016/j.ympev.2007.09.011

https://tree.opentreeoflife.org/curator/study/view/ot_1174?tab=trees&tree=tree3
Michael D. D'Emic, Roger B.J. Benson, 2013, 'Measurement, variation, and scaling of osteocyte lacunae: a case study in birds', Bone, vol. 57, no. 1, pp. 300-310
http://dx.doi.org/10.1016/j.bone.2013.08.010

https://tree.opentreeoflife.org/curator/study/view/pg_2860?tab=trees&tree=tree6646
Fulton, T. L., B. Letts, B. Shapiro. 2012. Multiple losses of flight and recent speciation in steamer ducks. Proceedings of the Royal Society B: Biological Sciences 279 (1737): 2339-2346.
http://dx.doi.org/10.1098/rspb.2011.2599

https://tree.opentreeoflife.org/curator/study/view/pg_2866?tab=trees&tree=tree6656
Gonzalez, J., H. Düttmann, M. Wink. 2009. Phylogenetic relationships based on two mitochondrial genes and hybridization patterns in Anatidae. Journal of Zoology 279 (3): 310-318.
http://dx.doi.org/10.1111/j.1469-7998.2009.00622.x

https://tree.opentreeoflife.org/curator/study/view/ot_1022?tab=trees&tree=tree1
Donne-Goussé, Carole, Vincent Laudet, Catherine Hänni. 2002. A molecular phylogeny of anseriformes based on mitochondrial DNA analysis. Molecular Phylogenetics and Evolution 23 (3): 339-356
http://dx.doi.org/10.1016/s1055-7903(02)00019-2

https://tree.opentreeoflife.org/curator/study/view/ot_1281?tab=trees&tree=Tr788
Rokas A., Melika G., Abe Y., Nieves-aldrey J., Cook J., & Stone G. 2003. Lifecycle closure, lineage sorting and hybridization revealed in a phylogenetic analysis of European oak gallwasps (Hymenoptera: Cynipidae: Cynipini) using mitochondrial sequence data. Molecular Phylogenetics and Evolution, 26(1): 36-45.
http://dx.doi.org/10.1016/S1055-7903(02)00329-9

https://tree.opentreeoflife.org/curator/study/view/pg_2858?tab=trees&tree=tree6642
Nylander, Johan A. A., Urban Olsson, Per Alström, Isabel Sanmartín. 2008. Accounting for phylogenetic uncertainty in biogeography: a Bayesian approach to dispersal-vicariance analysis of the thrushes (Aves: Turdus). Systematic Biology 57 (2): 257-268.
http://dx.doi.org/10.1080/10635150802044003

https://tree.opentreeoflife.org/curator/study/view/ot_159?tab=trees&tree=tree1
Benz, Brett W., Mark B. Robbins, A. Townsend Peterson. 2006. Evolutionary history of woodpeckers and allies (Aves: Picidae): placing key taxa on the phylogenetic tree. Molecular Phylogenetics and Evolution 40 (2): 389-399.
http://dx.doi.org/10.1016/j.ympev.2006.02.021

https://tree.opentreeoflife.org/curator/study/view/ot_178?tab=trees&tree=tree1
Sheldon, Frederick H., Linda A. Whittingham, Robert G. Moyle, Beth Slikas, David W. Winkler. 2005. Phylogeny of swallows (Aves: Hirundinidae) estimated from nuclear and mitochondrial DNA sequences. Molecular Phylogenetics and Evolution 35 (1): 254-270.
http://dx.doi.org/10.1016/j.ympev.2004.11.008

https://tree.opentreeoflife.org/curator/study/view/ot_314?tab=trees&tree=tree1
Suh, Alexander, Linnéa Smeds, Hans Ellegren. 2015. The dynamics of incomplete lineage sorting across the ancient adaptive radiation of neoavian birds. PLOS Biology13 (8): e1002224
http://dx.doi.org/10.1371/journal.pbio.1002224

https://tree.opentreeoflife.org/curator/study/view/ot_862?tab=trees&tree=tree1
Zhou, Xiaoping, Chengte Yao, Qingxian Lin, Wenzhen Fang, Xiaolin Chen. 2015. Complete mitochondrial genomes render the Night Heron genus Gorsachius non-monophyletic. Journal of Ornithology 157 (2): 505-513
http://dx.doi.org/10.1007/s10336-015-1297-z

https://tree.opentreeoflife.org/curator/study/view/ot_415?tab=trees&tree=tree2
García–R, Juan C., Gillian C. Gibb, Steve A. Trewick. 2014. Eocene diversification of crown group rails (Aves: Gruiformes: Rallidae). PLoS ONE 9 (10): e109635
http://dx.doi.org/10.1371/journal.pone.0109635

https://tree.opentreeoflife.org/curator/study/view/pg_1872?tab=trees&tree=tree3780
Fain, M., Krajewski C., & Houde P. 2007. Phylogeny of "core Gruiformes" (Aves: Grues) and resolution of the Limpkin-Sungrebe problem. Molecular Phylogenetics and Evolution 43 (2): 515-529.
http://dx.doi.org/10.1016/j.ympev.2007.02.015

https://tree.opentreeoflife.org/curator/study/view/ot_1490?tab=trees&tree=tree1
Nicholas G. Crawford, James F. Parham, Anna B. Sellas, Brant C. Faircloth, Travis C. Glenn, Theodore J. Papenfuss, James B. Henderson, Madison H. Hansen, W. Brian Simison, 2015, 'A phylogenomic analysis of turtles', Molecular Phylogenetics and Evolution, vol. 83, pp. 250-257
http://dx.doi.org/10.1016/j.ympev.2014.10.021

https://tree.opentreeoflife.org/curator/study/view/pg_1563?tab=trees&tree=tree6170
Sharkey, M., Carpenter J., Vilhelmsen L., Heraty J., Hawks D., Dowling A.P., Schulmeister S., Murray D., Deans A., Ronquist F., Krogmann L., & Wheeler W. 2011. Phylogenetic relationships among superfamilies of Hymenoptera. Cladistics 28 (1): 80-112.
http://dx.doi.org/10.1111/j.1096-0031.2011.00366.x

https://tree.opentreeoflife.org/curator/study/view/ot_750?tab=trees&tree=tree2
Zhi-Yong Yuan, Wei-Wei Zhou, Xin Chen, Nikolay A. Poyarkov, Hong-Man Chen, Nian-Hong Jang-Liaw, Wen-Hao Chou, Nicholas J. Matzke, Koji Iizuka, Mi-Sook Min, Sergius L. Kuzmin, Ya-Ping Zhang, David C. Cannatella, David M. Hillis, Jing Che, 2016, 'Spatiotemporal Diversification of the True Frogs (Genus Rana): A Historical Framework for a Widely Studied Group of Model Organisms', Systematic Biology, p. syw055
http://dx.doi.org/10.1093/sysbio/syw055

https://tree.opentreeoflife.org/curator/study/view/ot_146?tab=trees&tree=tree1
Baker, Allan J., Sergio L Pereira, Tara A Paton. 2007. Phylogenetic relationships and divergence times of Charadriiformes genera: multigene evidence for the Cretaceous origin of at least 14 clades of shorebirds. Biology. Letters 3 (2): 205-210.
http://dx.doi.org/10.1098/rsbl.2006.0606

https://tree.opentreeoflife.org/curator/study/view/pg_437?tab=trees&tree=tree6242
Wiegmann, B. M., M. D. Trautwein, I. S. Winkler, N. B. Barr, J.-W. Kim, C. Lambkin, M. A. Bertone, B. K. Cassel, K. M. Bayless, A. M. Heimberg, B. M. Wheeler, K. J. Peterson, T. Pape, B. J. Sinclair, J. H. Skevington, V. Blagoderov, J. Caravas, S. N. Kutty, U. Schmidt-Ott, G. E. Kampmeier, F. C. Thompson, D. A. Grimaldi, A. T. Beckenbach, G. W. Courtney, M. Friedrich, R. Meier, D. K. Yeates. 2011. Episodic radiations in the fly tree of life. Proceedings of the National Academy of Sciences. 108 (14): 5690-5695.
http://dx.doi.org/10.1073/pnas.1012675108

https://tree.opentreeoflife.org/curator/study/view/pg_2594?tab=trees&tree=tree6014
Lambkin, C., Sinclair B., Pape T., Courtney G.W., Skevington J., Meier R., Yeates D., Blagoderov V., & Wiegmann B.M. 2013. The phylogenetic relationships among infraorders and superfamilies of Diptera based on morphological evidence. Systematic Entomology 38 (1): 164-179.
http://dx.doi.org/10.1111/j.1365-3113.2012.00652.x

https://tree.opentreeoflife.org/curator/study/view/ot_179?tab=trees&tree=tree1
Lovette, Irby J., Brian S. Arbogast, Robert L. Curry, Robert M. Zink, Carlos A. Botero, John P. Sullivan, Amanda L. Talaba, Rebecca B. Harris, Dustin R. Rubenstein, Robert E. Ricklefs, Eldredge Bermingham. 2012. Phylogenetic relationships of the mockingbirds and thrashers (Aves: Mimidae). Molecular Phylogenetics and Evolution 63 (2): 219-229.
http://dx.doi.org/10.1016/j.ympev.2011.07.009

https://tree.opentreeoflife.org/curator/study/view/ot_180?tab=trees&tree=tree1
Lovette, Irby J., Dustin R. Rubenstein. 2007. A comprehensive molecular phylogeny of the starlings (Aves: Sturnidae) and mockingbirds (Aves: Mimidae): congruent mtDNA and nuclear trees for a cosmopolitan avian radiation. Molecular Phylogenetics and Evolution 44 (3): 1031-1056.
http://dx.doi.org/10.1016/j.ympev.2007.03.017

https://tree.opentreeoflife.org/curator/study/view/ot_771?tab=trees&tree=tree1
Martin Malmstrøm, Michael Matschiner, Ole K Tørresen, Bastiaan Star, Lars G Snipen, Thomas F Hansen, Helle T Baalsrud, Alexander J Nederbragt, Reinhold Hanel, Walter Salzburger, Nils C Stenseth, Kjetill S Jakobsen, Sissel Jentoft, 2016, 'Evolution of the immune system influences speciation rates in teleost fishes', Nature Genetics
http://dx.doi.org/10.1038/ng.3645

https://tree.opentreeoflife.org/curator/study/view/ot_1165?tab=trees&tree=tree1
Laura M. Zahn, 2017, 'Loss of flight in the Galapagos cormorant', Science, vol. 356, no. 6341, pp. 918.10-919
http://dx.doi.org/10.1126/science.356.6341.918-j

https://tree.opentreeoflife.org/curator/study/view/ot_140?tab=trees&tree=tree1
Kennedy, Martyn, Hamish G. Spencer. 2014. Classification of the cormorants of the world. Molecular Phylogenetics and Evolution 79: 249-257.
http://dx.doi.org/10.1016/j.ympev.2014.06.020

https://tree.opentreeoflife.org/curator/study/view/ot_1516?tab=trees&tree=Tr55113
Hu D., Xu X., Hou L., & Sullivan C. 2012. A new enantiornithine bird from the Lower Cretaceous of Western Liaoning, China, and its implications for early avian evolution. Journal of Vertebrate Paleontology, 32(3): 639-645.
http://dx.doi.org/10.1080/02724634.2012.652321

https://tree.opentreeoflife.org/curator/study/view/pg_2599?tab=trees&tree=tree6021
Alström, Per, Keith N. Barnes, Urban Olsson, F. Keith Barker, Paulette Bloomer, Aleem Ahmed Khan, Masood Ahmed Qureshi, Alban Guillaumet, Pierre-André Crochet, Peter G. Ryan. 2013. Multilocus phylogeny of the avian family Alaudidae (larks) reveals complex morphological evolution, non-monophyletic genera and hidden species diversity. Molecular Phylogenetics and Evolution 69 (3): 1043-1056.
http://dx.doi.org/10.1016/j.ympev.2013.06.005

https://tree.opentreeoflife.org/curator/study/view/ot_1185?tab=trees&tree=tree1
Chris J. Law, Graham J. Slater, Rita S. Mehta, 2017, 'Lineage Diversity and Size Disparity in Musteloidea: Testing Patterns of Adaptive Radiation Using Molecular and Fossil-Based Methods', Systematic Biology
http://dx.doi.org/10.1093/sysbio/syx047

https://tree.opentreeoflife.org/curator/study/view/ot_935?tab=trees&tree=tree2
Peter Rask Møller, Steen Wilhelm Knudsen, Werner Schwarzhans, Jørgen G. Nielsen, 2016, 'A new classification of viviparous brotulas (Bythitidae) – with family status for Dinematichthyidae – based on molecular, morphological and fossil data', Molecular Phylogenetics and Evolution, vol. 100, pp. 391-408
http://dx.doi.org/10.1016/j.ympev.2016.04.008

https://tree.opentreeoflife.org/curator/study/view/ot_863?tab=trees&tree=tree2
Scofield, R. Paul, Kieren J. Mitchell, Jamie R. Wood, Vanesa L. De Pietri, Scott Jarvie, Bastien Llamas, Alan Cooper. 2016. The Origin and Phylogenetic Relationships of the New Zealand Ravens. Molecular Phylogenetics and Evolution 106: 136–143
http://dx.doi.org/10.1016/j.ympev.2016.09.022

https://tree.opentreeoflife.org/curator/study/view/ot_289?tab=trees&tree=tree1
Dos Remedios, Natalie, Patricia L.M. Lee, Terry Burke, Tamás Székely, Clemens Küpper. 2015. North or south? Phylogenetic and biogeographic origins of a globally distributed avian clade. Molecular Phylogenetics and Evolution 89:151-159.
http://dx.doi.org/10.1016/j.ympev.2015.04.010

https://tree.opentreeoflife.org/curator/study/view/ot_176?tab=trees&tree=tree1
Cerasale, David J., Roi Dor, David W. Winkler, Irby J. Lovette. 2012. Phylogeny of the Tachycineta genus of New World swallows: insights from complete mitochondrial genomes. Molecular Phylogenetics and Evolution 63 (1): 64-71.
http://dx.doi.org/10.1016/j.ympev.2011.12.014

https://tree.opentreeoflife.org/curator/study/view/pg_1854?tab=trees&tree=tree3743
Mann, Nigel I., F. Keith Barker, Jeff A. Graves, Kimberly A. Dingess-Mann, Peter J.B. Slater. 2006. Molecular data delineate four genera of “Thryothorus” wrens. Molecular Phylogenetics and Evolution 40 (3): 750-759
http://dx.doi.org/10.1016/j.ympev.2006.04.014

https://tree.opentreeoflife.org/curator/study/view/ot_655?tab=trees&tree=tree1
Besnard, Guillaume, Joris A. M. Bertrand, Boris Delahaie, Yann X. C. Bourgeois, Emeline Lhuillier, Christophe Thébaud. 2015. Valuing museum specimens: high-throughput DNA sequencing on historical collections of New Guinea crowned pigeons (Goura). Biological Journal of the Linnean Society 117 (1): 71-82
http://dx.doi.org/10.1111/bij.12494

https://tree.opentreeoflife.org/curator/study/view/pg_2444?tab=trees&tree=tree6526
Pereira, S., Johnson K., Clayton D., and Baker A. 2007. Mitochondrial and nuclear DNA sequences support a Cretaceous origin of Columbiformes and a dispersal-driven radiation in the Paleogene. Systematic Biology 56 (4): 656-672.
http://dx.doi.org/10.1080/10635150701549672

https://tree.opentreeoflife.org/curator/study/view/ot_1846?tab=trees&tree=tree2
Wayne Maddison, Daiqin Li, Melissa Bodner, Junxia Zhang, Xu Xin, Qinqing Liu, Fengxiang Liu, 2014, 'The deep phylogeny of jumping spiders (Araneae, Salticidae)', ZooKeys, vol. 440, pp. 57-87
http://dx.doi.org/10.3897/zookeys.440.7891

https://tree.opentreeoflife.org/curator/study/view/ot_1638?tab=trees&tree=tree1
Bjarte H. Jordal, 2015, 'Molecular phylogeny and biogeography of the weevil subfamily Platypodinae reveals evolutionarily conserved range patterns', Molecular Phylogenetics and Evolution, vol. 92, pp. 294-307
http://dx.doi.org/10.1016/j.ympev.2015.05.028

https://tree.opentreeoflife.org/curator/study/view/pg_2870?tab=trees&tree=tree6662
Fuchs, Jérôme, Steven Chen, Jeff A. Johnson, David P. Mindell. 2011. Pliocene diversification within the South American forest falcons (Falconidae: Micrastur). Molecular Phylogenetics and Evolution 60 (3): 398-407.
http://dx.doi.org/10.1016/j.ympev.2011.05.008

https://tree.opentreeoflife.org/curator/study/view/pg_2871?tab=trees&tree=tree6663
Fuchs, Jérôme, Jeff A. Johnson, David P. Mindell. 2012. Molecular systematics of the caracaras and allies (Falconidae: Polyborinae) inferred from mitochondrial and nuclear sequence data. Ibis 154 (3): 520-532.
http://dx.doi.org/10.1111/j.1474-919X.2012.01222.x

https://tree.opentreeoflife.org/curator/study/view/ot_139?tab=trees&tree=tree1
Ramirez, J.L., C.Y. Miyaki, S.N. Del Lama. 2013. Molecular phylogeny of Threskiornithidae (Aves: Pelecaniformes) based on nuclear and mitochondrial DNA. Genetics and Molecular Research 12 (3): 2740-2750.
http://dx.doi.org/10.4238/2013.july.30.11

https://tree.opentreeoflife.org/curator/study/view/ot_768?tab=trees&tree=tree1
Slager, David L., C.J. Battey, Robert W. Bryson, Gary Voelker, John Klicka. 2014. A multilocus phylogeny of a major New World avian radiation: The Vireonidae. Molecular Phylogenetics and Evolution 80: 95-104
http://dx.doi.org/10.1016/j.ympev.2014.07.021

https://tree.opentreeoflife.org/curator/study/view/ot_1335?tab=trees&tree=Tr55332
Langer M.C., Bittencourt J.S., & Schultz C.L. 2011. A reassessment of the basal dinosaur Guaibasaurus candelariensis, from the Late Triassic Caturrita Formation of South Brazil. Earth and Environmental Science Transactions of the Royal Society of Edinburgh, 101: 301-332.
http://dx.doi.org/10.1017/S175569101102007X

https://tree.opentreeoflife.org/curator/study/view/ot_1449?tab=trees&tree=Tr55330
Shu-an J., Atterholt J., O'connor J.K., Lamanna M.C., Harris J.D., Da-qing L., Hai-lu Y., & Dodson P. 2011. A new, three-dimensionally preserved enantiornithine bird (Aves: Ornithothoraces) from Gansu Province, north-western China. Zoological Journal of the Linnean Society, 162(1): 201-219.
http://dx.doi.org/10.1111/j.1096-3642.2010.00671.x

https://tree.opentreeoflife.org/curator/study/view/ot_874?tab=trees&tree=tree1
Zuccon, Dario, Alice Cibois, Eric Pasquet, Per G.P. Ericson. 2006. Nuclear and mitochondrial sequence data reveal the major lineages of starlings, mynas and related taxa. Molecular Phylogenetics and Evolution 41 (2): 333-344
http://dx.doi.org/10.1016/j.ympev.2006.05.007

https://tree.opentreeoflife.org/curator/study/view/ot_172?tab=trees&tree=tree1
Päckert, Martin, Jochen Martens, Yue-Hua Sun, Lucia Liu Severinghaus, Alexander A. Nazarenko, Ji Ting, Till Töpfer, Dieter Thomas Tietze. 2011. Horizontal and elevational phylogeographic patterns of Himalayan and Southeast Asian forest passerines (Aves: Passeriformes). Journal of Biogeography 39 (3): 556-573.
http://dx.doi.org/10.1111/j.1365-2699.2011.02606.x

https://tree.opentreeoflife.org/curator/study/view/pg_1340?tab=trees&tree=tree2669
Hoy M., & Jeyaprakash A. 2008. First divergence time estimate of spiders, scorpions, mites and ticks (Subphylum: Chelicerata) inferred from mitochondrial phylogeny. Experimental and Applied Acarology, null.
http://dx.doi.org/10.1007/s10493-008-9203-5

https://tree.opentreeoflife.org/curator/study/view/ot_731?tab=trees&tree=tree2
Ottenburghs, Jente, Hendrik-Jan Megens, Robert H.S. Kraus, Ole Madsen, Pim van Hooft, Sipke E. van Wieren, Richard P.M.A. Crooijmans, Ronald C. Ydenberg, Martien A.M. Groenen, Herbert H.T. Prins. 2016. A Tree of Geese: A Phylogenomic Perspective on the Evolutionary History of True Geese. Molecular Phylogenetics and Evolution 101: 303-313.
http://dx.doi.org/10.1016/j.ympev.2016.05.021

https://tree.opentreeoflife.org/curator/study/view/pg_2691?tab=trees&tree=tree6244
Koepfli, Klaus-Peter, Matthew E. Gompper, Eduardo Eizirik, Cheuk-Chung Ho, Leif Linden, Jesus E. Maldonado, Robert K. Wayne. 2007. Phylogeny of the Procyonidae (Mammalia: Carnivora): Molecules, morphology and the Great American Interchange. Molecular Phylogenetics and Evolution 43 (3): 1076-1095.
http://dx.doi.org/10.1016/j.ympev.2006.10.003

https://tree.opentreeoflife.org/curator/study/view/ot_1023?tab=trees&tree=tree1
Peter A. Hosner, Edward L. Braun, Rebecca T. Kimball, 2015, 'Land connectivity changes and global cooling shaped the colonization history and diversification of New World quail (Aves: Galliformes: Odontophoridae)', Journal of Biogeography, vol. 42, no. 10, pp. 1883-1895
http://dx.doi.org/10.1111/jbi.12555

https://tree.opentreeoflife.org/curator/study/view/ot_158?tab=trees&tree=tree1
Moore, William S., Lowell C. Overton, Kathleen J. Miglia. 2011. Mitochondrial DNA based phylogeny of the woodpecker genera Colaptes and Piculus, and implications for the history of woodpecker diversification in South America. Molecular Phylogenetics and Evolution 58 (1): 76-84.
http://dx.doi.org/10.1016/j.ympev.2010.09.011
