## Supplemental Citations, Bird Families tree for "OpenTree: A Python package for Accessing and Analyzing data from the Open Tree of Life"

Citations for the phylogenies underlying the tree of bird families

https://tree.opentreeoflife.org/curator/study/view/ot_816?tab=trees&tree=tree1
Gibson, Rosemary, Allan Baker. 2012. Multiple gene sequences resolve phylogenetic relationships in the shorebird suborder Scolopaci (Aves: Charadriiformes). Molecular Phylogenetics and Evolution 64 (1): 66-72
http://dx.doi.org/10.1016/j.ympev.2012.03.008

https://tree.opentreeoflife.org/curator/study/view/ot_425?tab=trees&tree=tree1
Stein, R. Will, Joseph W. Brown, Arne Ø. Mooers. 2015. A molecular genetic time scale demonstrates Cretaceous origins and multiple diversification rate shifts within the order Galliformes (Aves). Molecular Phylogenetics and Evolution 92: 155-164
http://dx.doi.org/10.1016/j.ympev.2015.06.005

https://tree.opentreeoflife.org/curator/study/view/ot_809?tab=trees&tree=tree2
Jetz, W., G. H. Thomas, J. B. Joy, K. Hartmann, A. O. Mooers. 2012. The global diversity of birds in space and time. Nature 491 (7424): 444-448
http://dx.doi.org/10.1038/nature11631

https://tree.opentreeoflife.org/curator/study/view/ot_521?tab=trees&tree=tree1
Burleigh, J. Gordon, Rebecca T. Kimball, Edward L. Braun. 2015. Building the avian tree of life using a large-scale, sparse supermatrix. Molecular Phylogenetics and Evolution 84: 53-63
http://dx.doi.org/10.1016/j.ympev.2014.12.003

https://tree.opentreeoflife.org/curator/study/view/ot_156?tab=trees&tree=tree1
Moyle, Robert G. 2004. Phylogenetics of barbets (Aves: Piciformes) based on nuclear and mitochondrial DNA sequence data. Molecular Phylogenetics and Evolution 30 (1): 187-200.
http://dx.doi.org/10.1016/s1055-7903(03)00179-9

https://tree.opentreeoflife.org/curator/study/view/pg_2913?tab=trees&tree=tree6741
Price, Trevor D., Daniel M. Hooper, Caitlyn D. Buchanan, Ulf S. Johansson, D. Thomas Tietze, Per Alström, Urban Olsson, Mousumi Ghosh-Harihar, Farah Ishtiaq, Sandeep K. Gupta, Jochen Martens, Bettina Harr, Pratap Singh, Dhananjai Mohan. 2014. Niche filling slows the diversification of Himalayan songbirds. Nature 509: 222-225.
http://dx.doi.org/10.1038/nature13272

https://tree.opentreeoflife.org/curator/study/view/ot_520?tab=trees&tree=tree1
Jønsson, Knud Andreas, Pierre-Henri Fabre, Jonathan D. Kennedy, Ben G. Holt, Michael K. Borregaard, Carsten Rahbek, Jon Fjeldså. 2016. A supermatrix phylogeny of corvoid passerine birds (Aves: Corvides). Molecular Phylogenetics and Evolution 94: 87-94.
http://dx.doi.org/10.1016/j.ympev.2015.08.020

https://tree.opentreeoflife.org/curator/study/view/pg_2804?tab=trees&tree=tree6511
Kimball, Rebecca T., Colette M. St. Mary, Edward L. Braun. 2011. A macroevolutionary perspective on multiple sexual traits in the Phasianidae (Galliformes). International Journal of Evolutionary Biology 2011: 1-16.
http://dx.doi.org/10.4061/2011/423938

https://tree.opentreeoflife.org/curator/study/view/ot_312?tab=trees&tree=tree1
Schweizer, Manuel, Timothy F. Wright, Joshua V. Peñalba, Erin E. Schirtzinger, Leo Joseph. 2015. Molecular phylogenetics suggests a New Guinean origin and frequent episodes of founder-event speciation in the nectarivorous lories and lorikeets (Aves: Psittaciformes). Molecular Phylogenetics and Evolution 90: 34-48.
http://dx.doi.org/10.1016/j.ympev.2015.04.021

https://tree.opentreeoflife.org/curator/study/view/ot_171?tab=trees&tree=tree1
Spellman, GM, A Cibois, RG Moyle, K Winker, F Keith Barker. 2008. Clarifying the systematics of an enigmatic avian lineage: What is a bombycillid?. Molecular Phylogenetics and Evolution 49 (3): 1036-1040.
http://dx.doi.org/10.1016/j.ympev.2008.09.006

https://tree.opentreeoflife.org/curator/study/view/ot_290?tab=trees&tree=tree1
Selvatti, Alexandre Pedro, Luiz Pedreira Gonzaga, Claudia Augusta de Moraes Russo. 2015. A Paleogene origin for crown passerines and the diversification of the Oscines in the New World. Molecular Phylogenetics and Evolution 88: 1-15.
http://dx.doi.org/10.1016/j.ympev.2015.03.018

https://tree.opentreeoflife.org/curator/study/view/ot_783?tab=trees&tree=tree1
Moyle, Robert G., Carl H. Oliveros, Michael J. Andersen, Peter A. Hosner, Brett W. Benz, Joseph D. Manthey, Scott L. Travers, Rafe M. Brown, Brant C. Faircloth. 2016. Tectonic collision and uplift of Wallacea triggered the global songbird radiation. Nature Communications 7: 12709
http://dx.doi.org/10.1038/ncomms12709

https://tree.opentreeoflife.org/curator/study/view/pg_2015?tab=trees&tree=tree4152
Odeen, A., Hastad O., & Alstrom P. 2011. Evolution of ultraviolet vision in the largest avian radiation - the passerines. BMC Evolutionary Biology 11 (313): 1-8.
http://dx.doi.org/10.1186/1471-2148-11-313

https://tree.opentreeoflife.org/curator/study/view/ot_531?tab=trees&tree=tree1
Claramunt, Santiago, Joel Cracraft. 2015. A new time tree reveals Earth historys imprint on the evolution of modern birds. Science Advances 1 (11): e1501005-e1501005
http://dx.doi.org/10.1126/sciadv.1501005

https://tree.opentreeoflife.org/curator/study/view/ot_782?tab=trees&tree=Tr48736
Beresford, P., F.K. Barker, P.G. Ryan, T.M. Crowe. 2005. African endemics span the tree of songbirds (Passeri): molecular systematics of several evolutionary 'enigmas'. Proceedings of the Royal Society B: Biological Sciences 272 (1565): 849-858
http://dx.doi.org/10.1098/rspb.2004.2997

https://tree.opentreeoflife.org/curator/study/view/ot_1844?tab=trees&tree=tree1
Martin Stervander, Peter G. Ryan, Martim Melo, Bengt Hansson, 2019, 'The origin of the world’s smallest flightless bird, the Inaccessible Island Rail Atlantisia rogersi (Aves: Rallidae)', Molecular Phylogenetics and Evolution, vol. 130, pp. 92-98
http://dx.doi.org/10.1016/j.ympev.2018.10.007

https://tree.opentreeoflife.org/curator/study/view/pg_2575?tab=trees&tree=tree5974
Barker, F. K., K. J. Burns, J. Klicka, S. M. Lanyon, I. J. Lovette. 2013. Going to extremes: contrasting rates of diversification in a recent radiation of New World passerine birds. Systematic Biology 62 (2): 298-320.
http://dx.doi.org/10.1093/sysbio/sys094

https://tree.opentreeoflife.org/curator/study/view/ot_770?tab=trees&tree=tree1
Barker, F. Keith, Kevin J. Burns, John Klicka, Scott M. Lanyon, Irby J. Lovette. 2015. New insights into New World biogeography: An integrated view from the phylogeny of blackbirds, cardinals, sparrows, tanagers, warblers, and allies. The Auk 132 (2): 333-348.
http://dx.doi.org/10.1642/auk-14-110.1

https://tree.opentreeoflife.org/curator/study/view/pg_2924?tab=trees&tree=tree6755
Klicka, John, F. Keith Barker, Kevin J. Burns, Scott M. Lanyon, Irby J. Lovette, Jaime A. Chaves, Robert W. Bryson. 2014. A comprehensive multilocus assessment of sparrow (Aves: Passerellidae) relationships. Molecular Phylogenetics and Evolution 77: 177–182.
http://dx.doi.org/10.1016/j.ympev.2014.04.025

https://tree.opentreeoflife.org/curator/study/view/ot_841?tab=trees&tree=tree1
Simone Treplin, Romy Siegert, Christoph Bleidorn, Hazell Shokellu Thompson, Roger Fotso, Ralph Tiedemann, 2008, 'Molecular phylogeny of songbirds (Aves: Passeriformes) and the relative utility of common nuclear marker loci', Cladistics, vol. 24, no. 3, pp. 328-349
http://dx.doi.org/10.1111/j.1096-0031.2007.00178.x

https://tree.opentreeoflife.org/curator/study/view/pg_2913?tab=trees&tree=tree6742
Price, Trevor D., Daniel M. Hooper, Caitlyn D. Buchanan, Ulf S. Johansson, D. Thomas Tietze, Per Alström, Urban Olsson, Mousumi Ghosh-Harihar, Farah Ishtiaq, Sandeep K. Gupta, Jochen Martens, Bettina Harr, Pratap Singh, Dhananjai Mohan. 2014. Niche filling slows the diversification of Himalayan songbirds. Nature 509: 222-225.
http://dx.doi.org/10.1038/nature13272

https://tree.opentreeoflife.org/curator/study/view/ot_534?tab=trees&tree=tree1
Ksepka, Daniel T., Matthew J. Phillips. 2015. Avian diversification patterns across the K-Pg boundary: influence of calibrations, datasets, and model misspecification. Annals of the Missouri Botanical Garden 100 (4): 300-328
http://dx.doi.org/10.3417/2014032

https://tree.opentreeoflife.org/curator/study/view/ot_757?tab=trees&tree=Tr70438
Ericson, Per GP, Seraina Klopfstein, Martin Irestedt, Jacqueline MT Nguyen and Johan AA Nylander. 2014. Dating the diversification of the major lineages of Passeriformes (Aves). BMC Evolutionary Biology, 14(8).
http://dx.doi.org/10.1186/1471-2148-14-8

https://tree.opentreeoflife.org/curator/study/view/pg_420?tab=trees&tree=tree522
Hackett, S. J., R. T. Kimball, S. Reddy, R. C. K. Bowie, E. L. Braun, M. J. Braun, J. L. Chojnowski, W. A. Cox, K.-L. Han, J. Harshman, C. J. Huddleston, B. D. Marks, K. J. Miglia, W. S. Moore, F. H. Sheldon, D. W. Steadman, C. C. Witt, T. Yuri. 2008. A Phylogenomic Study of Birds Reveals Their Evolutionary History. Science 320 (5884): 1763-1768.
http://dx.doi.org/10.1126/science.1157704

https://tree.opentreeoflife.org/curator/study/view/ot_412?tab=trees&tree=Tr77157
Barker, F.K. 2014. Mitogenomic data resolve basal relationships among passeriform and passeridan birds. Molecular Phylogenetics and Evolution 79: 313-324.
http://dx.doi.org/10.1016/j.ympev.2014.06.011

https://tree.opentreeoflife.org/curator/study/view/ot_500?tab=trees&tree=tree1
Prum, Richard O., Jacob S. Berv, Alex Dornburg, Daniel J. Field, Jeffrey P. Townsend, Emily Moriarty Lemmon, Alan R. Lemmon. 2015. A comprehensive phylogeny of birds (Aves) using targeted next-generation DNA sequencing. Nature 526, (7574): 569-573
http://dx.doi.org/10.1038/nature15697

https://tree.opentreeoflife.org/curator/study/view/ot_532?tab=trees&tree=tree1
Gibb, Gillian C., Ryan England, Gerrit Hartig, P.A. (Trish) McLenachan, Briar L. Taylor Smith, Bennet J. McComish, Alan Cooper, David Penny. 2015. New Zealand passerines help clarify the diversification of major songbird lineages during the Oligocene. Genome Biology and Evolution 7 (11): 2983-2995.
http://dx.doi.org/10.1093/gbe/evv196

https://tree.opentreeoflife.org/curator/study/view/pg_2404?tab=trees&tree=tree5068
Barker, F. Keith, Alice Cibois, Peter Schikler, Julie Feinstein, Joel Cracraft. 2004. Phylogeny and diversification of the largest avian radiation. Proceedings of the National Academy of Sciences of the United States of America 101 (30): 11040-11045.
http://dx.doi.org/10.1073/pnas.0401892101

https://tree.opentreeoflife.org/curator/study/view/pg_2404?tab=trees&tree=tree6388
Barker, F. Keith, Alice Cibois, Peter Schikler, Julie Feinstein, Joel Cracraft. 2004. Phylogeny and diversification of the largest avian radiation. Proceedings of the National Academy of Sciences of the United States of America 101 (30): 11040-11045.
http://dx.doi.org/10.1073/pnas.0401892101

https://tree.opentreeoflife.org/curator/study/view/ot_854?tab=trees&tree=tree1
Ulf S. Johansson, Jon Fjeldså, Rauri C.K. Bowie, 2008, 'Phylogenetic relationships within Passerida (Aves: Passeriformes): A review and a new molecular phylogeny based on three nuclear intron markers', Molecular Phylogenetics and Evolution, vol. 48, no. 3, pp. 858-876
http://dx.doi.org/10.1016/j.ympev.2008.05.029

https://tree.opentreeoflife.org/curator/study/view/ot_111?tab=trees&tree=tree1
Alstrom, P., D. M. Hooper, Y. Liu, U. Olsson, D. Mohan, M. Gelang, H. Le Manh, J. Zhao, F. Lei, T. D. Price. 2014. Discovery of a relict lineage and monotypic family of passerine birds. Biology Letters 10 (3): 20131067-20131067.
http://dx.doi.org/10.1098/rsbl.2013.1067

https://tree.opentreeoflife.org/curator/study/view/pg_2872?tab=trees&tree=tree6665
Han, Kin-Lan, Mark B. Robbins, Michael J. Braun. 2010. A multi-gene estimate of phylogeny in the nightjars and nighthawks (Caprimulgidae). Molecular Phylogenetics and Evolution 55 (2): 443-453.
http://dx.doi.org/10.1016/j.ympev.2010.01.023

https://tree.opentreeoflife.org/curator/study/view/ot_753?tab=trees&tree=tree1
Wang, Ning, Rebecca T. Kimball, Edward L. Braun, Bin Liang, Zhengwang Zhang. 2016. Ancestral range reconstruction of Galliformes: the effects of topology and taxon sampling. Journal of Biogeography
http://dx.doi.org/10.1111/jbi.12782

https://tree.opentreeoflife.org/curator/study/view/pg_2577?tab=trees&tree=tree5980
Wang, Ning, Rebecca T. Kimball, Edward L. Braun, Bin Liang, Zhengwang Zhang. 2013. Assessing phylogenetic relationships among Galliformes: a multigene phylogeny with expanded taxon sampling in Phasianidae. PLoS ONE 8 (5): e64312.
http://dx.doi.org/10.1371/journal.pone.0064312

https://tree.opentreeoflife.org/curator/study/view/ot_1002?tab=trees&tree=tree1
Hosner, Peter A. Brant C. Faircloth, Travis C. Glenn, Edward L. Braun, Rebecca T. Kimball. 2015. Avoiding missing data biases in phylogenomic inference: an empirical study in the landfowl (Aves: Galliformes). Molecular Biology and Evolution 33 (4): 1110-1125
http://dx.doi.org/10.1093/molbev/msv347

https://tree.opentreeoflife.org/curator/study/view/ot_1345?tab=trees&tree=Tr109614
Worthy T., Mitri M., Handley W.D., Lee M.S., Anderson A., & Sand C. 2016. Osteology Supports a Stem-Galliform Affinity for the Giant Extinct Flightless Bird Sylviornis neocaledoniae (Sylviornithidae, Galloanseres). PLoS ONE, 11(3): e0150871.
http://dx.doi.org/10.1371/journal.pone.0150871

https://tree.opentreeoflife.org/curator/study/view/ot_116?tab=trees&tree=tree1
Hugall, Andrew F., Devi Stuart-Fox. 2012. Accelerated speciation in colour-polymorphic birds. Nature 485 (7400): 631-634.
http://dx.doi.org/10.1038/nature11050

https://tree.opentreeoflife.org/curator/study/view/ot_773?tab=trees&tree=tree1
Wood, Jamie R., Kieren J. Mitchell, R. Paul Scofield, Vanesa L. De Pietri, Nicolas J. Rawlence, Alan Cooper. 2016. Phylogenetic relationships and terrestrial adaptations of the extinct laughing owl, Sceloglaux albifacies (Aves: Strigidae). Zoological Journal of the Linnean Society
http://dx.doi.org/10.1111/zoj.12483

https://tree.opentreeoflife.org/curator/study/view/ot_837?tab=trees&tree=tree1
Robert G. Moyle, 2006, 'A MOLECULAR PHYLOGENY OF KINGFISHERS (ALCEDINIDAE) WITH INSIGHTS INTO EARLY BIOGEOGRAPHIC HISTORY', The Auk, vol. 123, no. 2, p. 487
http://dx.doi.org/10.1642/0004-8038(2006)123[487:ampoka]2.0.co;2

https://tree.opentreeoflife.org/curator/study/view/ot_855?tab=trees&tree=tree1
Martin Irestedt, Jérôme Fuchs, Knud A. Jønsson, Jan I. Ohlson, Eric Pasquet, Per G.P. Ericson, 2008, 'The systematic affinity of the enigmatic Lamprolia victoriae (Aves: Passeriformes)—An example of avian dispersal between New Guinea and Fiji over Miocene intermittent land bridges?', Molecular Phylogenetics and Evolution, vol. 48, no. 3, pp. 1218-1222
http://dx.doi.org/10.1016/j.ympev.2008.05.038

https://tree.opentreeoflife.org/curator/study/view/pg_2702?tab=trees&tree=tree6274
Aggerbeck, Marie, Jon Fjeldså, Les Christidis, Pierre-Henri Fabre, Knud Andreas Jønsson. 2014. Resolving deep lineage divergences in core corvoid passerine birds supports a proto-Papuan island origin. Molecular Phylogenetics and Evolution 70: 272-285.
http://dx.doi.org/10.1016/j.ympev.2013.09.027

https://tree.opentreeoflife.org/curator/study/view/ot_843?tab=trees&tree=tree3
Rheindt, Frank E., Janette A. Norman, Les Christidis. 2008. Phylogenetic relationships of tyrant-flycatchers (Aves: Tyrannidae), with an emphasis on the elaeniine assemblage. Molecular Phylogenetics and Evolution 46 (1): 88-101
http://dx.doi.org/10.1016/j.ympev.2007.09.011

https://tree.opentreeoflife.org/curator/study/view/ot_121?tab=trees&tree=tree7
Payne, Robert B. The cuckoos. Vol. 15. Oxford University Press, 2005. 
ISBN-13: 978-0198502135 ISBN-10: 0198502133
http://ukcatalogue.oup.com/product/9780198502135.do

https://tree.opentreeoflife.org/curator/study/view/ot_1174?tab=trees&tree=tree3
Michael D. D'Emic, Roger B.J. Benson, 2013, 'Measurement, variation, and scaling of osteocyte lacunae: a case study in birds', Bone, vol. 57, no. 1, pp. 300-310
http://dx.doi.org/10.1016/j.bone.2013.08.010

https://tree.opentreeoflife.org/curator/study/view/pg_2866?tab=trees&tree=tree6656
Gonzalez, J., H. Düttmann, M. Wink. 2009. Phylogenetic relationships based on two mitochondrial genes and hybridization patterns in Anatidae. Journal of Zoology 279 (3): 310-318.
http://dx.doi.org/10.1111/j.1469-7998.2009.00622.x

https://tree.opentreeoflife.org/curator/study/view/ot_314?tab=trees&tree=tree1
Suh, Alexander, Linnéa Smeds, Hans Ellegren. 2015. The dynamics of incomplete lineage sorting across the ancient adaptive radiation of neoavian birds. PLOS Biology13 (8): e1002224
http://dx.doi.org/10.1371/journal.pbio.1002224

https://tree.opentreeoflife.org/curator/study/view/pg_1586?tab=trees&tree=tree3208
Moyle, R.G., Andersen M.J., Oliveros C., Steinheimer F.D., and Reddy S. 2012. Phylogeny and biogeography of the core babblers (Aves: Timaliidae). Systematic Biology 61 (4): 631-651.
http://dx.doi.org/10.1093/sysbio/sys027

https://tree.opentreeoflife.org/curator/study/view/ot_415?tab=trees&tree=tree2
García–R, Juan C., Gillian C. Gibb, Steve A. Trewick. 2014. Eocene diversification of crown group rails (Aves: Gruiformes: Rallidae). PLoS ONE 9 (10): e109635
http://dx.doi.org/10.1371/journal.pone.0109635

https://tree.opentreeoflife.org/curator/study/view/pg_1872?tab=trees&tree=tree3780
Fain, M., Krajewski C., & Houde P. 2007. Phylogeny of "core Gruiformes" (Aves: Grues) and resolution of the Limpkin-Sungrebe problem. Molecular Phylogenetics and Evolution 43 (2): 515-529.
http://dx.doi.org/10.1016/j.ympev.2007.02.015

https://tree.opentreeoflife.org/curator/study/view/ot_836?tab=trees&tree=tree1
Robert G. Moyle, Joel Cracraft, Maklarin Lakim, Jamili Nais, Frederick H. Sheldon, 2006, 'Reconsideration of the phylogenetic relationships of the enigmatic Bornean Bristlehead (Pityriasis gymnocephala)', Molecular Phylogenetics and Evolution, vol. 39, no. 3, pp. 893-898
http://dx.doi.org/10.1016/j.ympev.2006.01.024

https://tree.opentreeoflife.org/curator/study/view/ot_847?tab=trees&tree=tree1
A. Manegold, 2008, 'Composition and phylogenetic affinities of vangas (Vangidae, Oscines, Passeriformes) based on morphological characters', Journal of Zoological Systematics and Evolutionary Research, vol. 46, no. 3, pp. 267-277
http://dx.doi.org/10.1111/j.1439-0469.2008.00458.x

https://tree.opentreeoflife.org/curator/study/view/pg_1953?tab=trees&tree=tree3977
Derryberry, E.P., Claramunt S., Derryberry G.E., Chesser R.T., Cracraft J., Aleixo A., Perez-eman J., Remsen J., and Brumfield R. 2011. Lineage diversification and morphological evolution in a large-scale continental radiation: the Neotropical ovenbirds and woodcreepers (Aves: Furnariidae). Evolution 65 (10): 2973-2986.
http://dx.doi.org/10.1111/j.1558-5646.2011.01374.x

https://tree.opentreeoflife.org/curator/study/view/ot_146?tab=trees&tree=tree1
Baker, Allan J., Sergio L Pereira, Tara A Paton. 2007. Phylogenetic relationships and divergence times of Charadriiformes genera: multigene evidence for the Cretaceous origin of at least 14 clades of shorebirds. Biology. Letters 3 (2): 205-210.
http://dx.doi.org/10.1098/rsbl.2006.0606

https://tree.opentreeoflife.org/curator/study/view/ot_180?tab=trees&tree=tree1
Lovette, Irby J., Dustin R. Rubenstein. 2007. A comprehensive molecular phylogeny of the starlings (Aves: Sturnidae) and mockingbirds (Aves: Mimidae): congruent mtDNA and nuclear trees for a cosmopolitan avian radiation. Molecular Phylogenetics and Evolution 44 (3): 1031-1056.
http://dx.doi.org/10.1016/j.ympev.2007.03.017

https://tree.opentreeoflife.org/curator/study/view/ot_1516?tab=trees&tree=Tr55113
Hu D., Xu X., Hou L., & Sullivan C. 2012. A new enantiornithine bird from the Lower Cretaceous of Western Liaoning, China, and its implications for early avian evolution. Journal of Vertebrate Paleontology, 32(3): 639-645.
http://dx.doi.org/10.1080/02724634.2012.652321

https://tree.opentreeoflife.org/curator/study/view/pg_2416?tab=trees&tree=tree3
Nunn, Gary B., Scott E. Stanley. 1998. Body size effects and rates of cytochrome b evolution in tube-nosed seabirds. Molecular Biology and Evolution 15 (10): 1360-1371.
http://dx.doi.org/10.1093/oxfordjournals.molbev.a025864

https://tree.opentreeoflife.org/curator/study/view/pg_2805?tab=trees&tree=tree6512
Wright, T. F., E. E. Schirtzinger, T. Matsumoto, J. R. Eberhard, G. R. Graves, J. J. Sanchez, S. Capelli, H. Muller, J. Scharpegge, G. K. Chambers, R. C. Fleischer. 2008. A multilocus molecular phylogeny of the parrots (Psittaciformes): support for a Gondwanan origin during the Cretaceous. Molecular Biology and Evolution 25 (10): 2141-2156.
http://dx.doi.org/10.1093/molbev/msn160

https://tree.opentreeoflife.org/curator/study/view/pg_2806?tab=trees&tree=tree6529
Joseph, Leo, Alicia Toon, Erin E. Schirtzinger, Timothy F. Wright. 2011. Molecular systematics of two enigmatic genera Psittacella and Pezoporus illuminate the ecological radiation of Australo-Papuan parrots (Aves: Psittaciformes). Molecular Phylogenetics and Evolution 59 (3): 675-684.
http://dx.doi.org/10.1016/j.ympev.2011.03.017

https://tree.opentreeoflife.org/curator/study/view/ot_655?tab=trees&tree=tree1
Besnard, Guillaume, Joris A. M. Bertrand, Boris Delahaie, Yann X. C. Bourgeois, Emeline Lhuillier, Christophe Thébaud. 2015. Valuing museum specimens: high-throughput DNA sequencing on historical collections of New Guinea crowned pigeons (Goura). Biological Journal of the Linnean Society 117 (1): 71-82
http://dx.doi.org/10.1111/bij.12494

https://tree.opentreeoflife.org/curator/study/view/pg_2444?tab=trees&tree=tree6526
Pereira, S., Johnson K., Clayton D., and Baker A. 2007. Mitochondrial and nuclear DNA sequences support a Cretaceous origin of Columbiformes and a dispersal-driven radiation in the Paleogene. Systematic Biology 56 (4): 656-672.
http://dx.doi.org/10.1080/10635150701549672

https://tree.opentreeoflife.org/curator/study/view/ot_874?tab=trees&tree=tree1
Zuccon, Dario, Alice Cibois, Eric Pasquet, Per G.P. Ericson. 2006. Nuclear and mitochondrial sequence data reveal the major lineages of starlings, mynas and related taxa. Molecular Phylogenetics and Evolution 41 (2): 333-344
http://dx.doi.org/10.1016/j.ympev.2006.05.007

https://tree.opentreeoflife.org/curator/study/view/pg_2869?tab=trees&tree=tree6661
Gonzalez, Juan-Carlos T., Ben C. Sheldon, Nigel J. Collar, Joseph A. Tobias. 2013. A comprehensive molecular phylogeny for the hornbills (Aves: Bucerotidae). Molecular Phylogenetics and Evolution 67 (2): 468-483.
http://dx.doi.org/10.1016/j.ympev.2013.02.012

https://tree.opentreeoflife.org/curator/study/view/pg_2926?tab=trees&tree=tree6757
Mitchell, K. J., B. Llamas, J. Soubrier, N. J. Rawlence, T. H. Worthy, J. Wood, M. S. Y. Lee, A. Cooper. 2014. Ancient DNA reveals elephant birds and kiwi are sister taxa and clarifies ratite bird evolution. Science 344 (6186): 898-900.
http://dx.doi.org/10.1126/science.1251981

https://tree.opentreeoflife.org/curator/study/view/ot_831?tab=trees&tree=tree2
ROBERT G. MOYLE, R TERRY CHESSER, RICHARD O. PRUM, PETER SCHIKLER, JOEL CRACRAFT, 2006, 'Phylogeny and Evolutionary History of Old World Suboscine Birds (Aves: Eurylaimides)', American Museum Novitates, vol. 3544, no. 1, p. 1
http://dx.doi.org/10.1206/0003-0082(2006)3544[1:paehoo]2.0.co;2
